# A Signature of Passivity? An Explorative Study of the N3 Event-Related Potential Component in Passive Oddball Tasks

**DOI:** 10.1101/271502

**Authors:** Boris Kotchoubey, Yuri G. Pavlov

## Abstract

**Background:** Many passive oddball experiments show a sharp negative deflection N3 after P3b, peaking between 400 and 500 ms, but this wave has never been analyzed properly. We conducted five passive oddball experiments, in which the number of deviants (i.e., one or two), their alleged meaning, and their distinctiveness varied.

**Results:** Mastoid- or common-referenced waveforms showed a fronto-central N3 in all experiments. The data were CSD (Current Source Density) transformed and underwent a Principal Component Analysis (PCA). The PCA revealed N3 containing two subcomponents with very stable peak latencies of about 415 and 455 ms, respectively. Both topography of the subcomponents and their variation with experimental conditions were very similar, indicating a midfrontal sink and a posterior temporal source. An analysis of P3a and P3b components replicated previously known effects.

**Conclusions:** We discuss the similarities and differences between the passive N3 and other components including the MMN, N1, late positive Slow Wave, and reorienting negativity. We also make general hypotheses about a possible functional meaning of N3; on this basis, specific hypotheses are formulated and further experiments are suggested to test these hypotheses.

## Introduction

More than half a century has passed since we first learned that infrequent stimuli, presented among highly frequent ones, elicit in the EEG a slow late positive wave with a typical peak latency slightly longer than 300 ms (Sutton et al., 1965). Already the first systematic review on this effect is almost four decades old (Donchin, 1981). Later on, this late positivity of the event-related potential (ERP), called P300 or P3, was found to entail two distinct components, P3a and P3b (e.g., Courchesne et al., 1975). Moreover, even P3b was shown to be not a unitary wave but consists of further subcomponents (e.g., Falkenstein et al., 1994).

The eliciting condition, referred to as the “oddball task” (Duncan-Johnson & Donchin, 1977) standardly involved an active task to respond to the rare stimuli or to both rare and frequent stimuli. If subjects have to respond to rare stimuli, the factors of rarity (or probability) and task-relevance are confounded. This fact underlay the idea to obtain the same effect without any active task requirement (Polich, 1987). It was shown that such a passive oddball paradigm also yields the P3 ERP component to the rare stimulus, although its amplitude is considerably smaller than in the active oddball paradigm (Lang et al., 1997; Polich, 1989). Although the exact component structure of this passive P3 remains unclear, the data indicate that it is not just a frontal P3a but contains at least a portion of the typical parieto-central P3b (e.g., Polich and McIsaak, 1994; Bennington and Pollich, 1999; Lang et al., 1997).

The interest to the passive P3 paradigm was particularly stimulated by clinical consideration, specifically by the idea to use cognitive ERP components for the analysis of cortical functions in behaviorally unresponsive patients, e.g., in vegetative state, minimally conscious state, in the late stages of neurodegenerative diseases, or in patients with Guillain-Barré syndrome. As these patients do not have any behavior, their cognitive abilities can only be examined by means of direct neurophysiological techniques, and the oddball ERP paradigm appears to be a convenient one. However, for the same reason of behavioral unresponsiveness the patients cannot be given an overt task – and if a task is covert (e.g., to count a rare stimulus), we cannot know whether they really perform it.

Several control studies aimed at the development of the passive oddball paradigm for severely brain damaged patients confirmed the presence of P3b, but the waveforms presented in these studies also indicated a later negative deflection with a peak latency between 400 and 500 ms. Bostanov & Kotchoubey (2006) used a continuous wavelet transform and clearly showed this wave in both time and time-frequency analyzed responses. Erlbeck et al. (2014) and Morlet et al. (2017) directly compared ERPs in the oddball paradigm (a) with an active instruction to respond to stimuli, (b) with active distraction of attention away from stimuli, and (c) with mind wandering (Morlet et al., 2017) or without any task (Erlbeck et al., 2014). Both studies showed a clear negativity following P3 to deviant stimuli in the last condition, but not in the other two. Unfortunately, the hypotheses these authors had developed for their oddball experiments concerned the Mismatch Negativity (MMN) and P3 components, thus the later components were not analyzed. A large, rather short-lasting, fronto-central negative wave after P3b can be seen clearly in the results obtained in passive oddball conditions with broadly various kinds of stimulation (e.g., Oades et al., 1995a, 1995b; Potts, 2004; Justen and Herbert, 2016). However, most of these studies were concentrated on P300 and/or MMN and did not analyze or even notice the post-P3 negativity. For example, O’Donnell et al. (1992) carefully described their ERPs as consisting of two negative-positive complexes N1-P2 and N2-P3 but ignored a clear negativity that followed P3; this was one of the first studies were such negativity can be seen in Figures.

Barry et al. (2006, 2009) were probably the first who explicitly mentioned this ERP component and termed it N3. However, the principal interest of this group was the relationship between the phase-locked ERPs and the underlying EEG activity. Therefore, even they did not perform a systematic analysis of N3 as compared with other ERP components even though they noticed some regularities of this component. Mueller et al. (2008) carefully investigated the life-span development of oddball ERP components (including N3) from early childhood to late adulthood. However, they did not clearly distinguish this rather sharp negative wave with a peak latency between 400-500 ms from the yet later, very slow component (peaking after 600 ms) that was also negative at frontal leads but inverted its polarity at parietal leads.

We found N3 accidently in Experiment I described below. This experiment, as well as some other studies showing a clear N3 (e.g., Justin & Herbert, 2016) contained an oddball related to learning and employed potentially significant stimuli. Therefore, we tested (i) the learning hypothesis in Experiment II (in which no learning took place altogether) and (ii) the significance hypothesis in Experiment IV (in which potentially significant stimuli were presented with increased salience) and in Experiment III (in which stimuli lost their significance). Also, because all these and many other similar experiments in the literature were three-stimulus oddballs, in Experiment V we asked (iii) whether a two-stimulus oddball would also yield N3.

The main aim of this explorative study was to show that N3 is not an occasional observation but a real and consistent ERP component, whose timing and topography are different from other known negativities recorded in oddball paradigms. Finally, we hypothesized that N3, being a late wave following the P3 complex, would demonstrate two important features common with this complex: firstly, N3 was expected to be larger to deviants than to standards; secondly, it was expected to decrease with repeated stimulation like P3 does (e.g., Polich, 1987; Polich and McIsaak, 1994).

To our best knowledge, the present study is the first ERP report specifically devoted to N3. For this reason, the study is largely exploratory. Of course, this is a serious limitation, which restraints the opportunities to design experiments on the basis of exact hypotheses. The reason is, however, that facts must be accumulated before precise hypotheses can be formulated.

## Methods: General

### Participants

Participants of all experiments were healthy individuals between 19 and 35 years, have had no present or prior diseases of the nervous system or hearing disorders. All of them reported being well aroused and in a good mood at the beginning of the experiment. None reported the use of any medicaments during the last weeks before the experiments. They were seated in comfortable chair and asked to close their eyes during EEG recording and to listen attentively to the stimuli presented through earphones. No other instruction was given.

The experiments were approved by the Ethical Commission of the University of Tübingen Medical Faculty. All participants gave their informed consent.

### Stimulation

Stimuli were chords each consisting of five harmonic frequencies and lasting for 200 ms. The intensity of all auditory stimuli was kept about 70 dB SPL. They were presented binaurally by means of pneumatic earphones (3M E-A-RTONE). All experiments contained a phase of passive oddball, in which one stimulus was presented frequently (Standard), and another one or two stimuli were rare deviants (Figure 1). Only the results of this phase (common to all experiments) will be reported. Stimuli were presented with SOA (onset-to-onset) varying between 950 and 1050 ms. The order of presentation was random except that one and the same deviant could not be delivered more than twice in a row.

**Figure 1.**
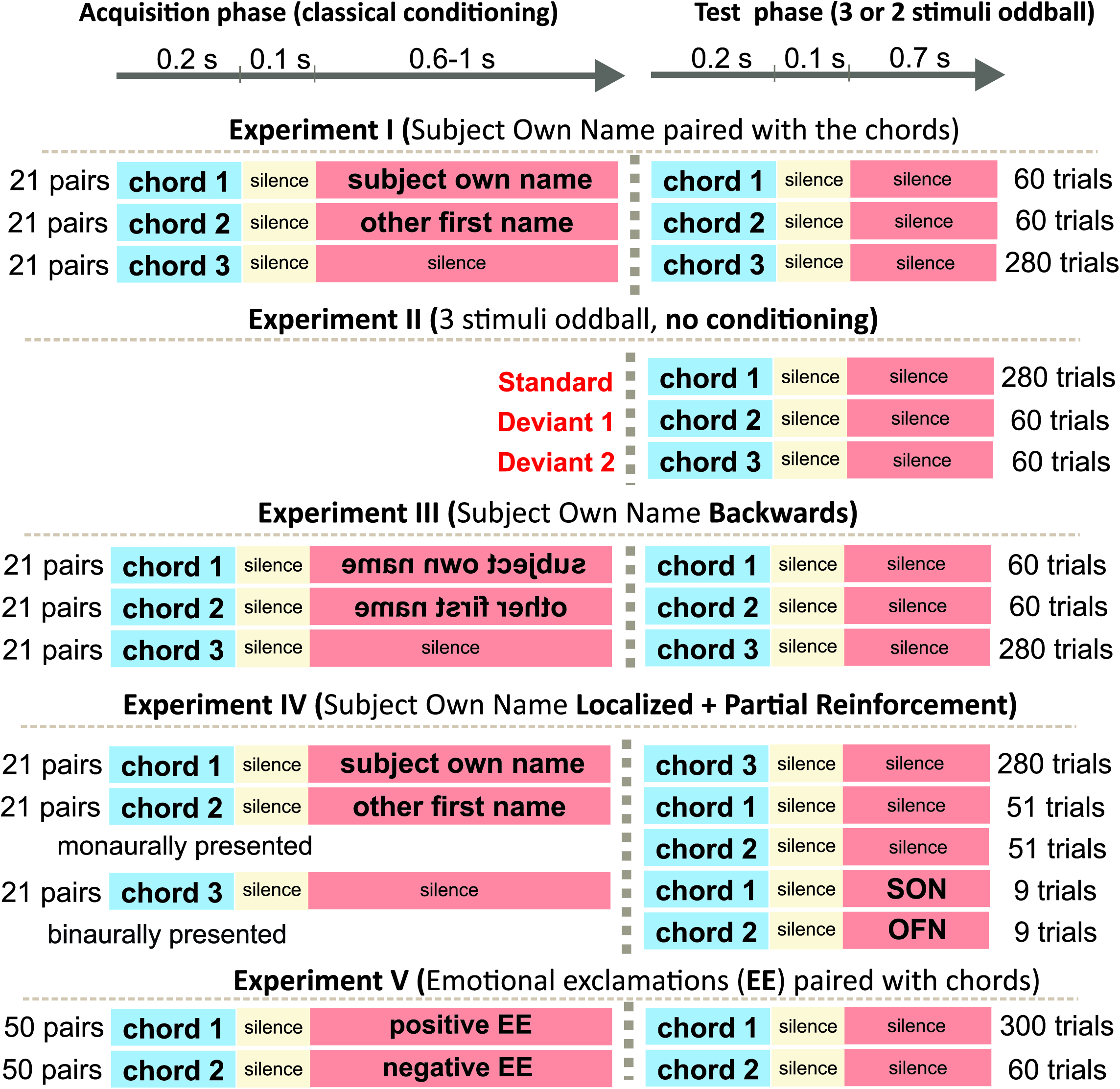
Experimental design.

### Recording

The EEG was recorded using 64 active ActiCHamp electrodes (Easycap GmbH, Herrsching, Germany) located according to the extended 10-20 system. In addition to the EEG, the vertical and horizontal electrooculograms were recorded. The impedance was below 25 kOhm. During recording, the EEG was referenced to Cz, the digitalization rate was 1000 Hz, and the low pass filter was 500 Hz.

### Preprocessing

The off-line inspection of the recordings revealed in some traces poor data quality in one or two of the 64 channels. After re-referencing to common average, these channels were replaced with interpolation of the adjacent electrodes. After this, the data were filtered within a band from 0.1 to 30 Hz. An Independent Component Analysis (ICA) was employed to separate and remove activity related to ocular artifacts according the AMICA algorithm (Palmer et al., 2012). The EEG was then broken into segments starting from 200 ms prior to tone onset and ending 800 ms after tone onset. The segments that still contained artifacts notwithstanding the preceding ICA correction were dismissed. The ERPs were averaged in relation to a baseline from −200 ms to 0 ms.

### Current Source Density (CSD)

We computed reference-free CSD estimates (μV/cm2 units; 10-cm head radius) by mean of the spherical spline surface Laplacian (Perrin et al., 1989) with the following parameters: 50 iterations; m = 4; smoothing constant λ = 10−5 (for detailed description of the procedure see Kayser and Tenke, 2006). CSD transformation helps to avoid problems associated with the choice of a recording reference. The CSD pattern of sources and sinks creates a clear image of the current generators of ERP components. This is especially important for auditory ERP when the use of linked-mastoids reference can lead to cancelling out potentials generated by sources in the temporal (auditory) cortex. Figure 2 shows a comparison between CSD and two other kinds of reference in Experiment II.

**Figure 2.**
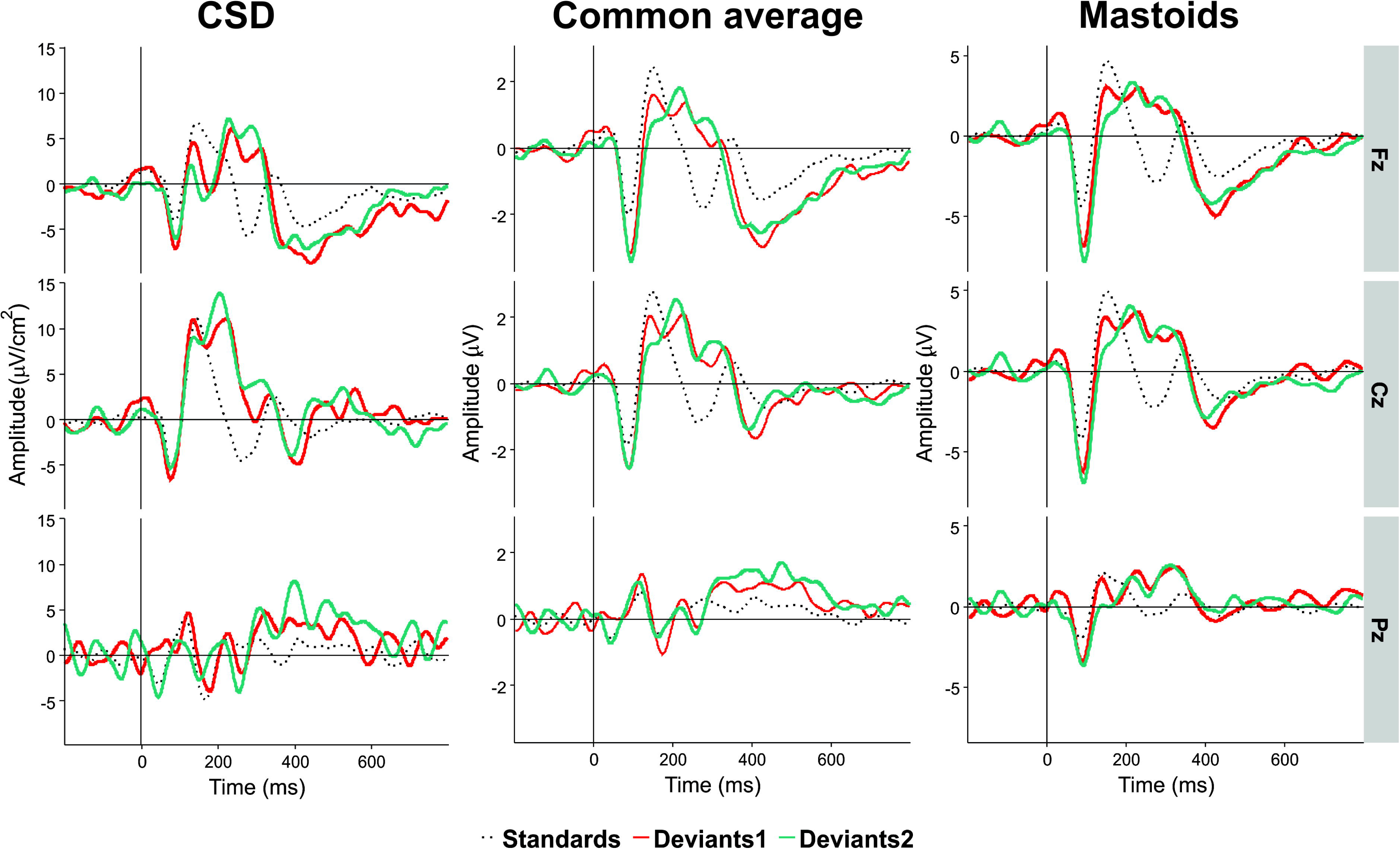
An illustration of the effect of electrical reference, for the results of Experiment II at three midline electrodes Fz, Cz, and Pz. Black dotted line, standards. Light red line, Deviant 1. Dark red line, Deviant 2. Positivity in this and the following figures is plotted upwards.

### Principal Component Analysis

After preprocessing, data were imported into the ERP PCA Toolkit (Dien, 2010). A temporal PCA with a Promax rotation was conducted to the combined dataset of CSD-ERPs averaged across trials where time intervals, conditions, electrodes sites and participants were considered as observations and time points as variables. In order to determine the number of factors to retain, the parallel test (Horn, 1965) was applied. The covariance matrix and Kaiser normalization were used for the PCA. The same procedure was applied, first, to a combined dataset of Experiment I to IV and then for Experiment V data separately. As a result of the parallel test, 55 temporal factors were extracted for rotation for Experiments I to IV and 44 for Experiment V. The waveforms for each factor were then reconstructed by multiplying factor scores by their corresponding loadings and standard deviations. CSD of reconstructed waveforms representing principal components of interest were averaged in 50 ms windows around peak values (±25 ms) in ROIs, time intervals and conditions. These data were used for statistical calculations.

The first eight principal components (PCs) with eigenvalues > 1 revealed close similarity with visually detected peaks (see Figure 3 for scalp distribution of the PCs). Particularly, PC1 with the peak latency of 93 ms was interpreted as N1; PC2 (412 ms), as an earlier peak of N3; PC3 (294 ms), as N2b; PC4 (456 ms), as a second subcomponent of N3; PC5 (215 ms), as P3a; PC6 (138 ms), as P2; and PC7 (334 ms), as P3b. Note that our N2b was recorded between P3a and P3b, while several studies reported N2b preceding P3a (e.g., Barry and De Blasio, 2015; Getzmann et al., 2015). Therefore, the reader may doubt whether it is the same component. We share this doubt but retain the name “N2b” because, first, no clear negative peak between P2 and P3b was manifested in our data; second, we were not interested in the true nature of N2b and analyzed this negativity only to compare it with N3.

**Figure 3.**
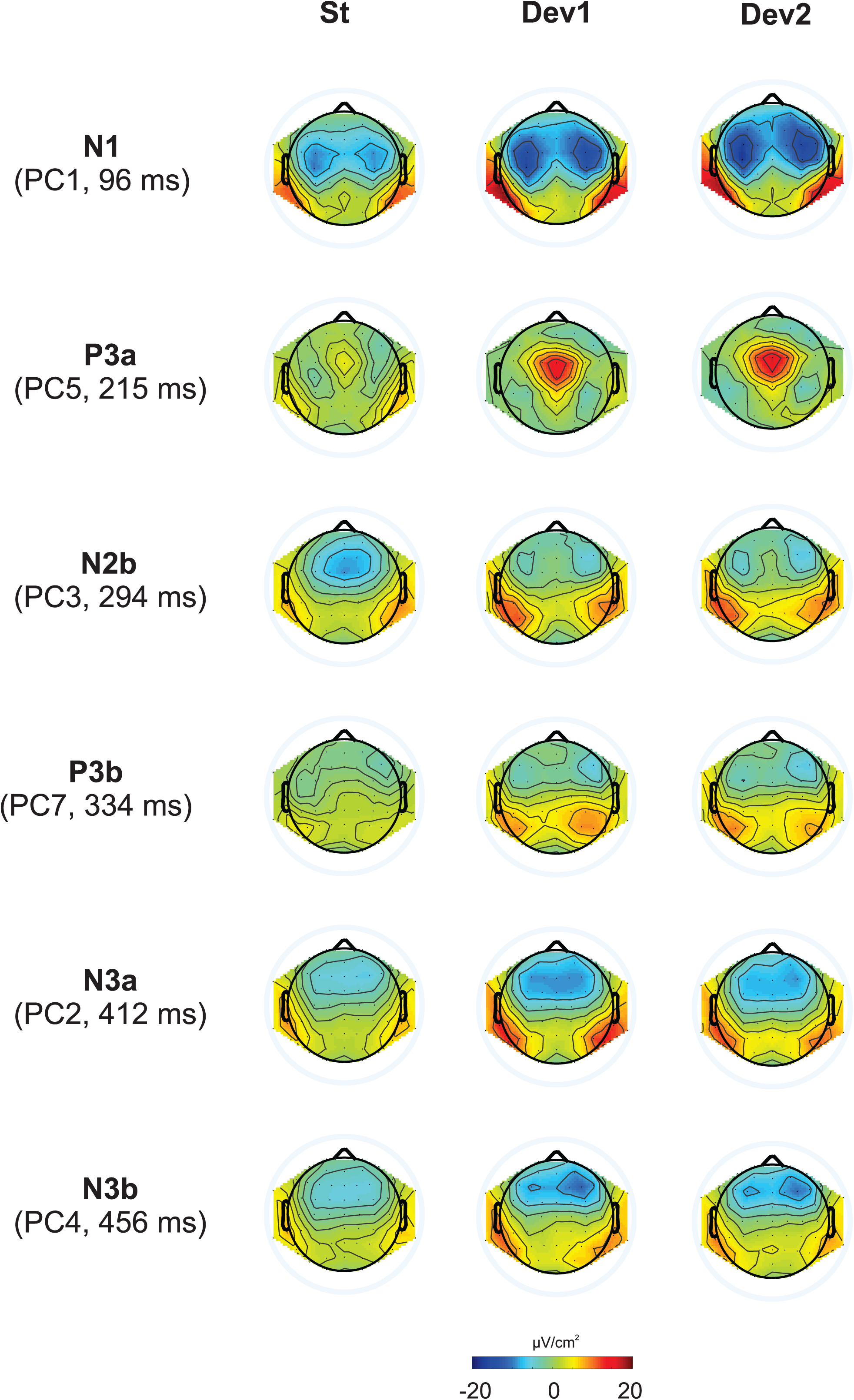
Topographic representation of each principal component and its peak latency, calculated for Current Source Density data. Because the results were nearly identical in Experiments I - IV, they are presented here averaged across the four three-stimuli experiments.

Because we had no prediction about the dynamics of P2, and to avoid unnecessary increase of the number of analyses, we did not analyze P2. Other components will be reported below with different number of details: (A) The N3 component will be reported in all details because, firstly, it is the main object of the study and, secondly, it has not been extensively studied in the literature, see above. (B) Other negative components N1 and N2b were analyzed not for their own sake but to compare them with N3. Finally, (C), the positive components P3a and P3b are, in contrast to N3, very well investigated in various kinds of oddball paradigm. Their analysis in the current study aimed solely at replication of already known effects.

### Statistical Analysis

As ERP components in oddball experiments are most typically investigated at the midline, our first analysis included only midline locations: frontal (mean of AFz, Fz and FCz), central (Cz), centroparietal (CPz) and parietal (Pz) leads. To explore the hypothesized changes with time, the responses were averaged separately for the first, second, and third thirds of the whole sequence of 400 stimuli. Each average included at least 18 (usually 20) deviants of each kind. The three levels of the factor Time will be referred to as T1 (stimuli 1 – 133), T2 (stimuli 134 – 266), and T3 (stimuli 267 – 400). The repeated-measures ANOVA included the factors Stimulus (3 levels), Site (4 levels), and Time (3 levels).

To analyze the activity over the hemispheres, five regions of interests (ROIs) were selected for each hemisphere: frontal (Fro), frontocentral (FC), central (Ce), posterior (Pos), and temporoparietal (TP), as shown in Figure 4. The corresponding repeated-measures ANOVA included the factors Stimulus (3 levels), Area (i.e., ROI, 5 levels), Hemisphere (2 levels), and Time (3 levels).

**Figure 4.**
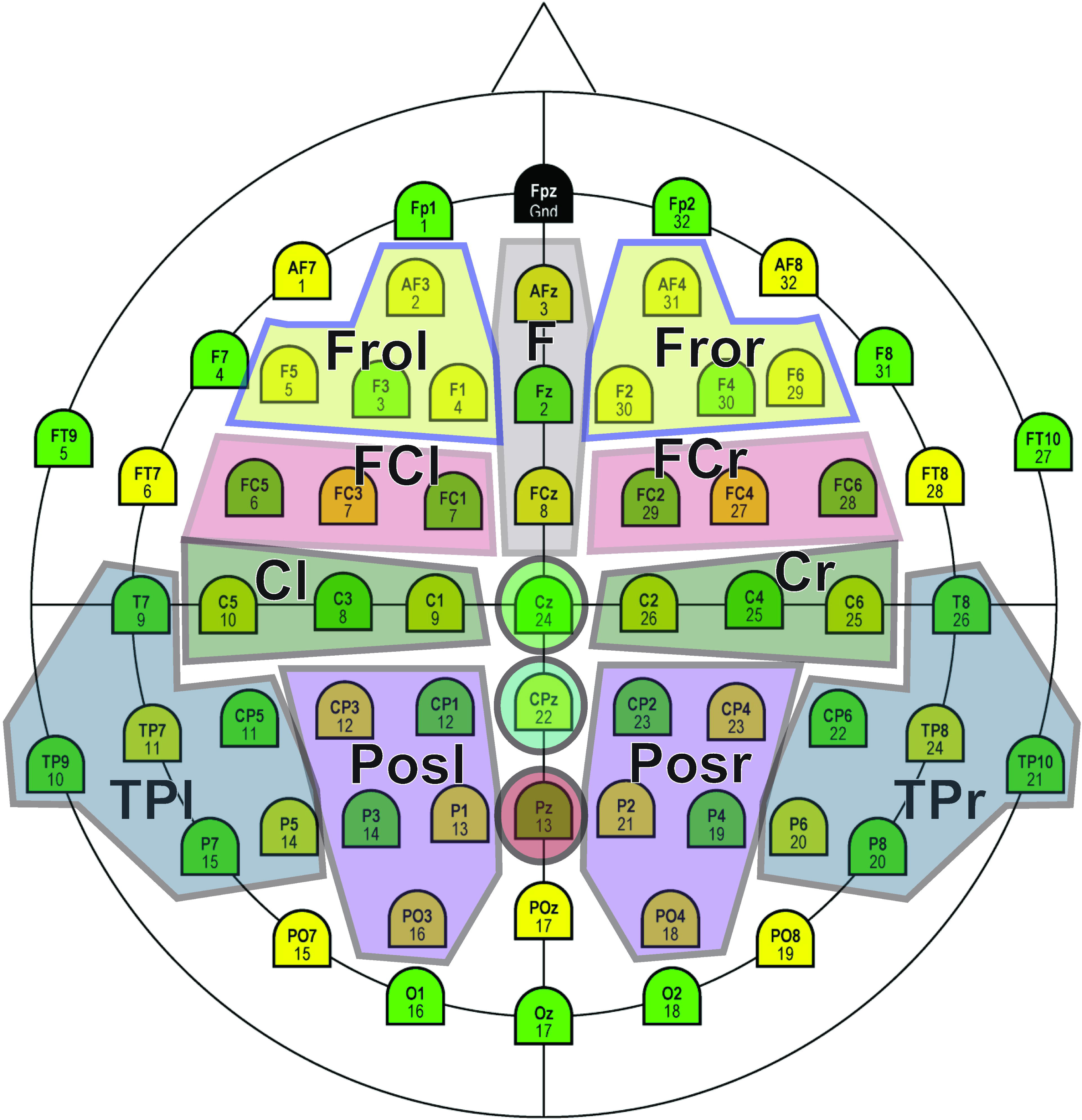
Regions of interest (ROIs) selected for statistical analysis.

The large total number of analyses might result in false positive effects (e.g., Luck & Gaspelin, 2017), and to date no exact method of correction exists. As a first approximation, we regarded the obtained Principal Components (PCs) as independent repetitions. Because in each experiment we analyzed six PCs, we shall generally report effects as “statistically significant” when p < 0.05/6 = 0.0083. However, the effects with .05 > p > .0083 will also be reported if they are repeatedly presented in similar conditions.

## Experiment I

### Methods

Twenty-three subjects (14 females) aged 19 to 33 (mean 25.2) participated in Experiment I. The experiment entailed two phases: Acquisition and Test. Chord 1, consisting of the frequencies 200, 400, 800, 1600, and 3200 Hz, was paired 21 times in the Acquisition phase in subjects with odd numbers with the own name of the corresponding participant (SON). Chord 2 consisted of 130, 260, 520, 1040, and 2080 Hz and was randomly paired 21 times with three other familiar names (OFN). Chord 3, consisting of 330, 660, 1320, 2640, and 5280 Hz, was presented 21 times without any relation to other stimuli. In subjects with even numbers, Chord 1 was paired with OFN, and Chord 2 with SON. The order of presentation was randomized but neither Chord 1 nor Chord 2 was allowed to appear more than 3 times in a row. The SOA within a pair chord-name was 300 ms. The interval after a pair was 1700-1800 ms, and the SOA after Chord 3 was 1150-1250 ms. Preliminary testing with eight healthy individuals showed that none of them had any difficulty to distinguish between the tones.

The average duration of the own name and the other names was 669 ms (SD = 9 ms) and 676 ms (SD = 12 ms) respectively (t = 0.78, p = .44). Other names originated from the same pool of the most frequent German names used for each subject’s own name, and always contained the same number of syllables as the own name.

In the Test phase (passive oddball) Chord 1 was presented 280 times, and Chord 2 and Chord 3, 60 times each. Name stimuli were not presented any longer. In other words, Test phase was a typical three-stimulus oddball paradigm with Chord 3 as a standard and Chords 1 and 2 as deviants.

### Results

Figure 5 presents the overview of the obtained ERP waveforms. Table 1 shows the main statistical results across components and experiments for midline recording sites. The ANOVA results for lateral electrode sites are shown in Table 2, and the statistics obtained in each ROI are summarized in Table 3. In the following text only sporadic statistical results are explained that are not resumed in the Tables.

**Figure 5.**
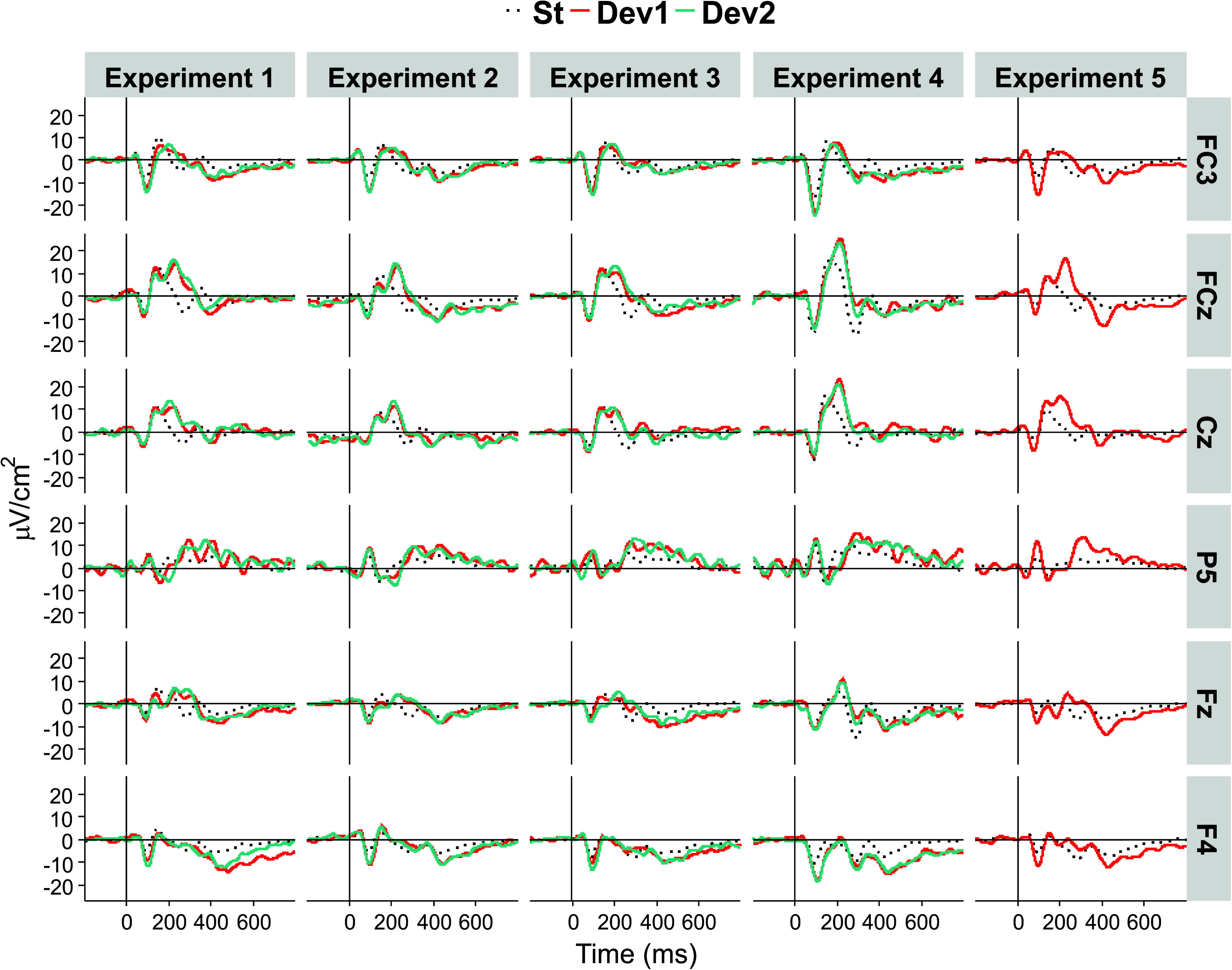
Average CSD ERP waveforms in each of the five experiments, at selected locations.

**Table 1.** Results of ANOVA Site x Stimulus x Time in midline electrodes

|  | Experiment I | Experiment II | Experiment III | Experiment IV | Experiment V |
| --- | --- | --- | --- | --- | --- |
| **N3 early** |  |  |  |  |  |
| Site | F(3,66) = 5.77, p = .015, $\eta^2 = .21$ | F(3,57) = 17.73, p < .001, $\eta^2 = .48$ | F(3,60) = 21.87, p < .001, $\eta^2 = .52$ | F(3,63) = 20.45, p < .001, $\eta^2 = .49$ | F(3,63) = 42.94, p < .001, $\eta^2 = .67$ |
| Stimulus | | F(2,38) = 6.53, p = .004, $\eta^2 = .26$ | | | |
| Stimulus:Site | | F(6,114) = 2.85, p = .039, $\eta^2 = .13$ | F(6,120) = 6.55, p = .001, $\eta^2 = .25$ | | F(3,63) = 15.39, p < .001, $\eta^2 = .42$ |
| Stimulus:Time | | | | F(6,126) = 4.38, p = .003, $\eta^2 = .17$ | |
| **N3 late** |  |  |  |  |  |
| Site | F(2,44) = 14.53, p < .001, $\eta^2 = .42$ | F(3,57) = 14.82, p < .001, $\eta^2 = .44$ | F(3,60) = 17.76, p < .001, $\eta^2 = .47$ | F(3,63) = 36.25, p < .001, $\eta^2 = .63$ | F(3,63) = 44.29, p < .001, $\eta^2 = .68$ |
| Stimulus:Site | | F(6,114) = 2.94, p = .025, $\eta^2 = .12$ | F(6,120) = 3.20, p = .026, $\eta^2 = .15$ | | F(3,63) = 5.98, p = .004, $\eta^2 = .22$ |
| **N1** |  |  |  |  |  |
| Site | F(3,66) = 6.70, p = .006, $\eta^2 = .23$ | F(3,57) = 5.99, p = .006, $\eta^2 = .24$ | F(3,60) = 8.54, p = .004, $\eta^2 = .30$ | F(3,63) = 34.17, p < .001, $\eta^2 = .62$ | F(3,63) = 18.50, p < .001, $\eta^2 = .47$ |
| Stimulus | | | F(2,40) = 5.87, p = .006, $\eta^2 = .23$ | | |
| Stimulus:Site | | | | | F(3,63) = 7.04, p = .003, $\eta^2 = .25$ |
| **N2b** |  |  |  |  |  |
| Site | F(3,66) = 3.89, p = .028, $\eta^2 = .15$ | | F(3,60) = 4.98, p = .010, $\eta^2 = .20$ | F(3,63) = 19.86, p < .001, $\eta^2 = .46$ | F(3,63) = 9.73, p = .001, $\eta^2 = .32$ |
| Stimulus | | F(2,38) = 21.63, p < .001, $\eta^2 = .53$ | F(2,40) = 5.79, p = .009, $\eta^2 = .23$ | F(2,42) = 8.98, p = .001, $\eta^2 = .30$ | F(1,21) = 19.21, p < .001, $\eta^2 = .48$ |
| Stimulus:Site | | | | F(6,126) = 6.03, p < .001, $\eta^2 = .22$ | |
| Stimulus:Time | F(4,88) = 4.27, p = .008, $\eta^2 = .16$ | | | | |

| P3a |  |  |  |  |  |
| --- | --- | --- | --- | --- | --- |
| Site | F(2,44) = 9.57,<br>p < .001,<br>$\eta^2 = .30$ | F(3,57) = 4.53,<br>p = .025,<br>$\eta^2 = .19$ | | | |
| Stimulus | F(2,44) = 15.83,<br>p < .001,<br>$\eta^2 = .42$ | F(2,38) = 8.86,<br>p = .002,<br>$\eta^2 = .32$ | F(2,40) = 20.22,<br>p < .001,<br>$\eta^2 = .50$ | F(2,42) = 37.80,<br>p < .001,<br>$\eta^2 = .60$ | F(1,21) = 23.00,<br>p < .001,<br>$\eta^2 = .52$ |
| Time | F(2,44) = 3.88,<br>p = .049,<br>$\eta^2 = .15$ | | F(2,40) = 5.52,<br>p = .009,<br>$\eta^2 = .22$ | F(2,42) = 10.55,<br>p < .001,<br>$\eta^2 = .33$ | F(2,42) = 3.55,<br>p = .039,<br>$\eta^2 = .15$ |
| Stimulus:Site | F(6,132) = 3.77,<br>p = .020,<br>$\eta^2 = .15$ | F(6,114) = 6.06,<br>p = .004,<br>$\eta^2 = .24$ | F(6,120) = 5.17,<br>p = .001,<br>$\eta^2 = .21$ | F(6,126) = 9.39,<br>p < .001,<br>$\eta^2 = .31$ | F(3,63) = 13.01,<br>p < .001,<br>$\eta^2 = .39$ |
| P3b |  |  |  |  |  |
| Site | | | F(3,60) = 9.44,<br>p < .001,<br>$\eta^2 = .33$ | F(3,63) = 9.81,<br>p = .001,<br>$\eta^2 = .32$ | |
| Stimulus | | | | | F(1,21) = 9.12,<br>p = .007,<br>$\eta^2 = .30$ |
| Time | | | | | F(2,42) = 4.23,<br>p = .022,<br>$\eta^2 = .17$ |
| Stimulus:Time | | | | F(4,84) = 3.16,<br>p = .033,<br>$\eta^2 = .13$ | |
| Stimulus:Site | F(6,132) = 3.36,<br>p = .020,<br>$\eta^2 = .13$ | | F(6,120) = 2.92,<br>p = .037,<br>$\eta^2 = .13$ | | F(3,63) = 3.34,<br>p = .045,<br>$\eta^2 = .14$ |

**Table 2.** Results of ANOVA Area x Hemisphere x Stimulus x Time in lateral electrodes

|  | Experiment I | Experiment II | Experiment III | Experiment IV | Experiment V |
| --- | --- | --- | --- | --- | --- |
| **N3 early** |  |  |  |  |  |
| Area | F(4,88) = 36.26,<br>p < .001,<br>$\eta^2 = .62$ | F(4,76) = 48.48,<br>p < .001,<br>$\eta^2 = .72$ | F(4,80) = 45.48,<br>p < .001,<br>$\eta^2 = .70$ | F(4,84) = 39.54,<br>p < .001,<br>$\eta^2 = .65$ | F(4,84) = 66.25,<br>p < .001,<br>$\eta^2 = .76$ |
| Stimulus | | F(2,38) = 4.56,<br>p = .018,<br>$\eta^2 = .19$ | | F(2,42) = 4.41,<br>p = .018,<br>$\eta^2 = .17$ | |
| Stimulus:Area | F(8,176) = 5.34,<br>p = .004,<br>$\eta^2 = .20$ | F(8,152) = 5.90,<br>p < .001,<br>$\eta^2 = .24$ | F(8,160) = 6.55,<br>p = .001,<br>$\eta^2 = .25$ | F(8,168) = 5.09,<br>p = .001,<br>$\eta^2 = .19$ | F(4,84) = 21.88,<br>p < .001,<br>$\eta^2 = .51$ |
| **N3 late** |  |  |  |  |  |
| Area | F(4,88) = 48.25,<br>p < .001,<br>$\eta^2 = .69$ | F(4,76) = 27.34,<br>p < .001,<br>$\eta^2 = .59$ | F(4,80) = 28.32,<br>p < .001,<br>$\eta^2 = .59$ | F(4,84) = 45.84,<br>p < .001,<br>$\eta^2 = .69$ | F(4,84) = 59.92,<br>p < .001,<br>$\eta^2 = .74$ |
| Stimulus:Area | F(8,176) = 7.08,<br>p < .001,<br>$\eta^2 = .24$ | F(8,152) = 3.54,<br>p = .005,<br>$\eta^2 = .16$ | F(8,160) = 4.12,<br>p = .003,<br>$\eta^2 = .17$ | F(8,168) = 5.81,<br>p = .001,<br>$\eta^2 = .22$ | F(4,84) = 7.24,<br>p < .001,<br>$\eta^2 = .26$ |
| **N1** |  |  |  |  |  |
| Area | F(4,88) = 38.78,<br>p < .001,<br>$\eta^2 = .64$ | F(4,76) = 28.85,<br>p < .001,<br>$\eta^2 = .60$ | F(4,80) = 24.65,<br>p < .001,<br>$\eta^2 = .55$ | F(4,84) = 59.46, p<br>< .001,<br>$\eta^2 = .74$ | F(4,84) = 50.00,<br>p < .001,<br>$\eta^2 = .70$ |
| Stimulus | F(2,44) = 33.72,<br>p < .001,<br>$\eta^2 = .61$ | F(2,38) = 69.35,<br>p < .001,<br>$\eta^2 = .79$ | F(2,40) = 31.20,<br>p < .001,<br>$\eta^2 = .61$ | F(2,42) = 28.25, p<br>< .001,<br>$\eta^2 = .64$ | F(1,21) = 73.22,<br>p < .001,<br>$\eta^2 = .78$ |
| Stimulus: Area | F(8,176) =<br>11.48, p < .001,<br>$\eta^2 = .34$ | F(8,152) = 9.78,<br>p < .001,<br>$\eta^2 = .34$ | F(8,160) = 10.07,<br>p < .001,<br>$\eta^2 = .34$ | F(8,168) = 6.48, p<br>< .001,<br>$\eta^2 = .24$ | F(4,84) = 34.15,<br>p < .001,<br>$\eta^2 = .62$ |
| Area:Hemisphere | F(4,88) = 4.53,<br>p = .007,<br>$\eta^2 = .17$ | | | | |
| **N2b** |  |  |  |  |  |
| Area | F(4,88) = 27.79,<br>p < .001,<br>$\eta^2 = .56$ | F(4,76) = 10.12,<br>p = .001,<br>$\eta^2 = .35$ | F(4,80) = 17.38,<br>p < .001,<br>$\eta^2 = .46$ | F(4,84) = 37.84,<br>p < .001,<br>$\eta^2 = .64$ | F(4,84) = 50.31,<br>p < .001,<br>$\eta^2 = .71$ |
| Stimulus | F(2,44) = 20.90,<br>p < .001,<br>$\eta^2 = .49$ | F(2,38) = 24.50,<br>p < .001,<br>$\eta^2 = .56$ | F(2,40) = 27.21,<br>p < .001,<br>$\eta^2 = .58$ | F(2,42) = 6.18,<br>p = .005,<br>$\eta^2 = .22$ | F(1,21) = 45.47,<br>p < .001,<br>$\eta^2 = .68$ |
| **P3a** |  |  |  |  |  |
| Area | | | | | F(4,84) = 6.02,<br>p = .005,<br>$\eta^2 = .22$ |
| Stimulus | F(2,44) = 6.43,<br>p = .008,<br>$\eta^2 = .23$ | | | | |
| Area:Hemisphere | F(4,76) = 3.33,<br>p = .029,<br>$\eta^2 = .15$ | | F(4,80) = 4.68,<br>p = .008,<br>$\eta^2 = .19$ | F(4,84) = 3.31,<br>p = .026,<br>$\eta^2 = .14$ | |
| Stimulus: Area | F(8,146) = 14.13, p < .001,<br>$\eta^2 = .39$ | F(8,152) = 6.28,<br>p < .001,<br>$\eta^2 = .25$ | F(8,160) = 6.70,<br>p < .001,<br>$\eta^2 = .25$ | F(8,168) = 14.06,<br>p < .001,<br>$\eta^2 = .40$ | F(4,84) = 9.13,<br>p < .001,<br>$\eta^2 = .30$ |
| **P3b** |  |  |  |  |  |
| Area | F(4,88) = 17.77,<br>p < .001,<br>$\eta^2 = .45$ | F(4,76) = 6.82,<br>p = .001,<br>$\eta^2 = .26$ | F(4,80) = 16.58,<br>p < .001,<br>$\eta^2 = .45$ | F(4,84) = 19.55,<br>p < .001,<br>$\eta^2 = .48$ | F(4,84) = 13.97,<br>p < .001,<br>$\eta^2 = .38$ |
| Stimulus | F(2,44) = 12.04,<br>p < .001,<br>$\eta^2 = .35$ | F(2,38) = 5.99,<br>p = .011,<br>$\eta^2 = .24$ | F(2,40) = 10.18,<br>p = .001,<br>$\eta^2 = .34$ | F(2,42) = 3.47,<br>p = .048,<br>$\eta^2 = .14$ | F(1,21) = 35.33,<br>p < .001,<br>$\eta^2 = .63$ |
| Time | F(2,44) = 9.56,<br>p < .001,<br>$\eta^2 = .30$ | | F(2,40) = 3.97,<br>p = .042,<br>$\eta^2 = .17$ | F(2,42) = 4.58,<br>p = .021,<br>$\eta^2 = .18$ | |
| Time:Area | F(8,176) = 3.12,<br>p = .013,<br>$\eta^2 = .12$ | F(8,152) = 3.55,<br>p = .023,<br>$\eta^2 = .16$ | | | |
| Stimulus: Area | F(8,176) = 5.76,<br>p = .001,<br>$\eta^2 = .21$ | F(8,152) = 3.75,<br>p = .009,<br>$\eta^2 = .16$ | F(8,160) = 3.11,<br>p = .018,<br>$\eta^2 = .13$ | F(8,168) = 6.88,<br>p < .001,<br>$\eta^2 = .25$ | F(4,84) = 6.57,<br>p = .003,<br>$\eta^2 = .24$ |

**Table 3.** Analysis of Area x Stimulus interactions for negative ERP components at lateral sites: ANOVA results

| Area | Experiment I | Experiment II | Experiment III | Experiment IV | Experiment V |
| --- | --- | --- | --- | --- | --- |
| **N3 early** |  |  |  |  |  |
| Fro<br>D— | | F(2,38) = 4.34, p =<br>.021, $\eta^2$ = .22; | F(2,40) = 3.99, p =<br>.036, $\eta^2$ = .14 | F(2,42) = 9.87, p <<br>.001, $\eta^2$ = .32 | F(1,21) = 25.47, p <<br>.001, $\eta^2$ = .52 |
| FC<br>D— | F(2,44) = 10.49, p <<br>.001, $\eta^2$ = .32 | F(2,38) = 4.47, p =<br>.030, $\eta^2$ = .23 | F(2,40) = 4.51, p =<br>.018, $\eta^2$ = .18 | F(2,42) = 11.59, p <<br>.001, $\eta^2$ = .36 | F(1,21) = 28.63, p <<br>.001, $\eta^2$ = .58 |
| Ce<br>D— |  |  |  | . |  |
| TP<br>D+ | | (F(2,38) = 4.31, p =<br>.020, $\eta^2$ = .18 | F(2,40) = 3.74, p =<br>.045, $\eta^2$ = .16 | | F(1,21) = 11.71, p =<br>.003, $\eta^2$ = .36 |
| **N3 late** |  |  |  |  |  |
| Fro<br>D— | F(2,44) = 11.23, p <<br>.001, $\eta^2$ = .34 | F(2,38) = 3.40, p =<br>.011, $\eta^2$ = .18 | F(2,40) = 8.24, p =<br>.003, $\eta^2$ = .48 | F(2,42) = 12.40,<br>p < .001, $\eta^2$ = .37 | F(1,21) = 10.09, p =<br>.005, $\eta^2$ = .32 |
| FC<br>D— | F(2,44) = 8.11, p =<br>.001, $\eta^2$ = .27 | F(2,38) = 13.32, p <<br>.001, $\eta^2$ = .41 | | F(2,42) = 5.90, p =<br>.006, $\eta^2$ = .22 | F(1,21) = 5.27, p =<br>.032, $\eta^2$ = .20 |
| Ce<br>D— | F(2,44) = 6.51, p =<br>.004, $\eta^2$ = .23 (§) | F(2,38) = 6.70, p =<br>.003, $\eta^2$ = .26 | | | |
| TP<br>D+ | F(2,44) = 9.43, p =<br>.001, $\eta^2$ = .30 | F(2,38) = 10.87, p <<br>.001, $\eta^2$ = .26 | | . | |
| **N1** |  |  |  |  |  |
| Fro<br>D— | F(2,44) = 16.53, p <<br>.001, $\eta^2$ = .43 | F(2,38) = 16.45, p <<br>.001, $\eta^2$ = .46 | F(2,40) = 7.44, p =<br>.013, $\eta^2$ = .27 | F(2,42) = 19.21, p <<br>.001, $\eta^2$ = .48 | |
| FC<br>D— | F(2,44) = 20.95, p <<br>.001, $\eta^2$ = .49 | F(2,38) = 37.82, p <<br>.001, $\eta^2$ = .67 | | F(2,42) = 13.42, p<br>< .001, $\eta^2$ = .39 | F(1,21) = 11.73, p =<br>.003, $\eta^2$ = .36 |
| Ce<br>D— | F(2,44) = 31.19, p <<br>.001, $\eta^2$ = .59 | F(2,38) = 20.00, p <<br>.001, $\eta^2$ = .51 | F(2,40) = 8.38, p =<br>.001, $\eta^2$ = .30 (*) | F(2,42) = 11.03, p <<br>.001, $\eta^2$ = .34 | F(1,21) = 52.45, p <<br>.001, $\eta^2$ = .69 |
| TP<br>D+ | F(2,44) = 11.27, p <<br>.001, $\eta^2$ = .35 | F(2,38) = 9.63, p <<br>.001, $\eta^2$ = .34 | | F(2,42) = 4.80, p =<br>.021, $\eta^2$ = .19 | F(1,21) = 13.42, p =<br>.001, $\eta^2$ = .39 |

The table presents main effects of Stimulus at single areas when the Stimulus x Area interaction was significant. Fro, frontal areas; FC, fronto-central areas; Ce, central areas; TP, lateral temporoparietal areas. Posterior areas are omitted because no effect attained $p \leq .05$ there. D— means that the amplitude was more negative to deviants than standards. D+ means that the amplitude was more positive to deviants than to standards. Empty cells mean that, although Stimulus x Area interactions were significant, the effect of Stimulus in a given area was not.
(§) Chord 2 more negative than Chord 1: $t(22) = 2.90$ , $p = .008$ , $\eta^2 = .28$
(\*) This was a Stimulus x Time interaction with deviants being significantly more negative than standards at time intervals T2 and T3

N3, the main object of the present study, revealed in the PCA as entailing two principal components, **PC2** and **PC4**, with peak latencies of 412 and 456 ms, respectively. Both PCs were negative at frontal sites and positive at Pz, resulting in a main effect of Site. A similar anterior-posterior negativity-positivity slope was also significant in lateral sites. In addition, strong Stimulus x Area interactions indicated that this slope was much stronger for deviants than standards, thus demonstrating an oddball effect (see Table 3).

The component **N1 (PC1)** had a frontal maximum at the midline (main effect of Site, see Table 1). Lateral amplitudes were characterized by strong regional differences with positive values over TP areas and negative values in all other areas (main effect of Area) and more negative amplitudes in response to deviants than standards (main effect of Stimulus). This latter general effect was, however, inverted over TP areas where the amplitudes were more positive to deviants than standards, resulting in significant Stimulus x Area interactions.

Over posterior areas, where no stimulus effect was found, the amplitudes were more negative on the left than right side (t(22) = 3.56, p < .001, η^2^ = .47). The opposite holds true for TP areas (t(22) = 2.91, p = .008, η^2^ = .40), yielding a significant Area x Hemisphere interaction.

**N2b (PC3)** was characterized by positive values in response to deviants but negative to standards, resulting in a main effect of Stimulus at midline. This effect was modified by Time, as Chord 2 elicited positive PC3/N2b values at T3 like the standard. Also at the lateral sites the same effects of Stimulus (i.e., positivity to deviants, negativity to standards) and topographic differences (i.e., significantly positive amplitudes in the Pos and TP areas, and slightly negative amplitudes in the Fro and FC areas) were found.

**P3a** (**PC5)** was larger to deviants than standards (main Stimulus effect). This difference had its maximum at Cz and minimum at Pz (Stimulus x Site interaction). Moreover, the amplitude was generally largest at Cz (main effect of Site). Further, the amplitude of P3a habituated from T1 to T2 and T3 (i.e., main effect of Time).

Over the lateral areas P3a to standards increased in the posterior direction and reached its maximum in TP areas. The amplitude to deviants was, in contrast, maximal in FC areas and minimal in TP areas, resulting in a strong Stimulus x Area interaction (see Table 2). In general, the amplitude was more positive to deviants than standards (main effect of Stimulus). In TP areas the amplitude (particularly in response to standards) was more positive on the right than left side, which manifested itself in the interaction between Area and Hemisphere.

The amplitude of **P3b (PC7)** was larger to deviants than standards at Pz (Stimulus x Site interaction in the midline ANOVA) and at the lateral sites (main effect of Stimulus in the lateral ANOVA). The maximal positive deviant-standard difference was found in Pos and TP areas, whereas in Fro and FC areas the amplitudes were even more negative to deviants than standards. Anterior negativities were larger on the right than left side (Area x Hemisphere interaction: F(4,88) = 3.15, p = .036, η^2^ = .13).

### Discussion

In addition to the expected oddball effects on P3a and P3b, the experiment showed a robust N3 wave following P3b. Also this wave was characterized by a strong oddball effect, and its spatial distribution was different from other negativities (N1, N2b).

The experiment employed a passive oddball paradigm with potentially significant (previously associated with SON and OFN) stimuli. An obvious hypothesis (which could also be supported by similar data from the literature, see Introduction), was that N3 is related to the significance of deviants or to the learning process preceding oddball. On this basis the following three experiments were designed. Specifically, Experiment III was designed to reduce the significance, and Experiment IV was, in contrast, expected to enhance the salience of the stimuli (thus enhancing the ERP effects). Experiment II was expected to eliminate preceding learning.

## Experiment II

### Methods

Experiments II, III, and IV were carried out with the same participants, always in the same order. The resulting samples were not identical, however. Specifically, the instruction to close eyes resulted in the almost complete absence of ocular activity. On the other hand, two participants exhibited signs of sleep stages N1 or even N2 (AASM sleep criteria) in their EEG, and their data had to be dismissed. The data of seven males and 13 females, aged 22 to 29 (mean 25.6) were analyzed. None of the participants of Experiments II to IV participated in Experiments I or V.

The design was fully identical to the Test phase of Experiment I. However, no Acquisition phase preceded it. The only instruction was to keep the eyes closed and to listen to stimuli.

### Results

In addition to the Site effect like in Experiment I, the amplitude of both **N3** subcomponents was more negative to deviants than standards at frontal sites (F(2,40) = 6.46, p = .004, η^2^ = .24, 3.95, p = .032, η^2^ = .17 for PC2 and PC4, respectively), and more positive to deviants than standards at Pz (F(2,40) = 4.51, p = .023, η^2^ = .18, and 3.33, p = .051, η^2^ = .15 for PC2 and PC4, respectively). At the lateral areas, strong effects of Area and Stimulus x Area interactions indicated the same differential effects of standards and deviants as in Experiment I above. The exact statistical results are shown in Tables 1 – 3.

The effects on **N1** were almost identical to those in Experiment I. Regarding **N2b**, additionally to the effects like those in Experiment I, lateral frontal and fronto-central negativities significantly decreased with Time (main effect of Time: F(4,76) = 10.12, p = .001, η^2^ = .35).

Also the effects on the positive components were similar to Experiment I with the following differences: the Area x Hemisphere interaction for P3a was not significant; the oddball effect at the midline for P3b (i.e., the Stimulus x Site interaction) was not significant. The Area x Hemisphere interaction for P3b was, again, significant: F(4,76) = 5.00, p = .004, η^2^ = .21.

## Experiment III

### Methods

After exclusion of one data set for technical reasons, data of twenty-one subjects (12 females, aged 22 – 29, mean 25.7) were analyzed.

The design was identical to Experiment I including both Acquisition and Test phases. However, the words presented during Acquisition were made completely unrecognizable at the same time preserving all auditory features that SON and OFN had in Experiment I. Note that a simple reversion of the audio track would not be enough for this sake because correctly pronounced and inverted names have quite different dependence of intensity on time. Therefore, a more complex masking procedure was employed. First, the first 25% of time points of an original name were multiplied by a linearly spaced vector of coefficients from 1.5 to 0, and the remaining 75% points were set to 0. Second, the first 25% of time points of the same name played backwards were multiplied by a linearly spaced vector of coefficients from 0 to 1.5, and the last 75% time points remained unchanged. Third, the two files were added. The processing was done in Matlab. A pilot experiment with 40 medical students (not participating in the EEG experiments) revealed than none of them was able to recognize the presented words, including each participant’ masked own name.

### Results

The effects on **N3** at midline were similar to Experiment I. PC2 was again more negative to deviants than standards at frontal sites (F(2,40) = 6.46, p = .004, η^2^ = .24), and this effect was inverted at Pz (F(2,40) = 4.51, p = .023, η^2^ = .18). PC4 was more negative to deviants than standards at Cz (F(2,40) = 4.11, p = .024, η^2^ = .17) with a similar non-significant frontal tendency (F(2,40) = 3.12, p = .057, η^2^ = .14) and the lack of Stimulus effect at CPz and Pz. Lateral effects were identical for PC2 and PC4, i.e., the amplitudes were significantly negative (and more negative to deviants than standards) over the Fro and FC areas, and significantly positive (and more positive to deviants than standards) over the TP areas (see Tables 2 and 3).

The amplitude of **N1** was significantly larger to Chord 1 (previously linked with masked SON) than Chord 2 (previously linked with masked OFN), while the standard stimulus elicited an intermediate amplitude between the two deviants. At the lateral sites, deviants elicited larger negativities in frontal areas (F(1,20 = 7.44, p = .013, η^2^ = .27) and in central areas at T2 and T3 (Stimulus x Time interaction with F(2,40) = 8.38, p = .001, η^2^ = .30). The standard-deviant differences in other areas were not significant.

The effects on **N2b** were quite like Experiment II, including the decrease of lateral negativities with Time (F(2,40) = 4.48, p = .020, η^2^ = .18) that was lacking in Experiment I. The effects on **P3a** and **P3b** were similar to Experiment I including the larger anterior negativities on the right than left side (Area x Hemisphere interaction: F(4,80) = 3.70, p = .016, η^2^ = .16).

### Discussion

Contrary to our hypothesis, changes in N3 in Experiments II and III were almost the same as in Experiment I although the significance of the eliciting stimuli substantially decreased. In Experiment III the tones had previously been associated not with participants’ own names and other familiar names (as in Experiment I) but with unrecognizable acoustical sequences. In Experiment II the tones were completely neutral. Their significance was so low that two of the 22 participants felt asleep, and the others complained of boredom. In accordance with these observations, participants’ reports after Experiments II and III were not exact, as many of them reported having heard “two or three” different sounds. Even though the same effects in Experiment II were, generally, weaker than in Experiment I, this fact can most plausibly be attributed to the lower alertness but not to some specific effects of learning.

The idea of Experiment IV was, in contrast to Experiments II and III, to increase the (possibly insufficient) salience of auditory stimuli. This was done by means of (i) monaural presentation, and (ii), partial reinforcement. In addition, we wanted to rule out the supposition that the effects might be related with the particular set of stimuli used in Experiments I to III.

## Experiment IV

### Methods

Participants were 13 females and 9 males, aged 22 – 29, mean 25.7. The design was similar to Experiment I with three differences. First, different tones were used. Chord 1 (associated with SON) contained the harmonic components of 300, 600, 120, 2400, and 4800 Hz; chord 2 (associated with OFN) contained the harmonics of 195, 390, 780, 1560, and 3120 Hz; and the standard contained the components of 495, 990, 1980, 3960, and 7920 Hz. Second, Chords 1 and 2 were presented monaurally, one to the left ear and the other one to the right ear, with the side of presentation being counterbalanced among participants. Third, the Test phase included partial reinforcement. The tone related to SON in the Acquisition phase was also followed by SON on randomly selected nine (of the 60) trials in the Test phase; on the remaining 51 trials the tone was presented alone like in Experiment I, Likewise, the tone related to OFN in the Acquisition phase was also followed by OFN on 9 trials in the Test phase. The partial reinforcement aimed at the refreshment of the association between tones and names.

### Results

Only the main Site effect on **N3** at midline replicated the corresponding effect in the previous experiments, but Stimulus x Site interactions did not attain the corrected significance level of .0083. PC2 revealed, in addition, a significant Site x Time interaction, indicating the increasing positivity at Pz at T3 as compared with T1 and T2 (main Time effect at Pz: F(2,42) = 3.46, p = .038, η^2^ = .15). Interestingly, the Stimulus x Site interaction for PC2 was significant when only two deviant stimuli were compared (F(3,63) = 6.09, p = .006, η^2^ = .26) indicating a significantly more negative frontal amplitude to the OFN-associated tone than to the SON-associated tone (t(21) = 2.57, p = .018, η^2^ = .24).

In the lateral areas the effects of Area and Stimulus x Area interactions were similar to the other experiments, as can be seen in Tables 2 and 3. In addition to the data of the Tables, the OFN-associated tone elicited a larger FC negativity (main effect of Stimulus for PC2: F(1,21) = 7.22, p = .014, η^2^ = .26) and a larger TP positivity at T1 (Stimulus x Time interaction: F(2,42) = 3.62, p = .036, η^2^ = .15) than the SON-associated tone. Note, however, that these effects were “significant” only with the *uncorrected* significance level.

The effects on **N1** were the same as in Experiment I. In addition to already depicted effects on **N2b**, a significant Stimulus X Site interaction (F(6,126) = 6.03, p < .001, η^2^ = .22) indicated that the anterior-posterior inversion from negative to positive polarity took place for deviants at more anterior sites than for standards.

The effects on **P3a** were identical to those in Experiments I and III, including the habituation of the component (i.e., main effect of Time). There was no oddball effect on **P3b** at Pz (like Experiment II, but differing from Experiments I and III). In contrast, the significant Stimulus x Time interaction was due to the fact that at T2 and T3 the deviant previously linked with OFN elicited larger responses than the previously linked with SON (main effect of Stimulus at T3: F(2,42) = 5.74, p = .007, η^2^ = .22, with the mean amplitudes of 1.47, 2.97, and 0.26 µV/cm^2^ to standard, the former deviant, and the latter deviant, respectively).

### Discussion

At a behavioral level, the manipulation yielded the expected improvement of participants’ reports as compared with the preceding experiments. All participants correctly reported having heard three different tones, two of which were associated with their own names or with other familiar names. ERP results were, nevertheless, quite similar to the results of all previous experiments. The following two analyses were intended to check this similarity.

## Three experiments

### Methods

Nineteen participants (11 females, aged 22-29, mean 25.5±0.48) yielded analyzable data in all three Experiments (II, III and IV). Therefore, we performed additional ANOVAs with the results of these subjects for the three experiments together. The analyses included the same repeated measures factors plus the additional factor Experiment (3 levels).

### Results

The analysis of **N3** revealed only one interaction that approached (but not reached) the corrected significance level of .0083. The interaction resulted from topographic differences for PC4 being larger in Experiment IV than in Experiments I and III (Experiment x Area interaction for lateral electrodes: F(8,144) = 3.43, p = .009, η^2^ = .16). Figure 6 illustrates the comparative dynamics of different ERP components at the sites of their maximal expression.

**Figure 6.**
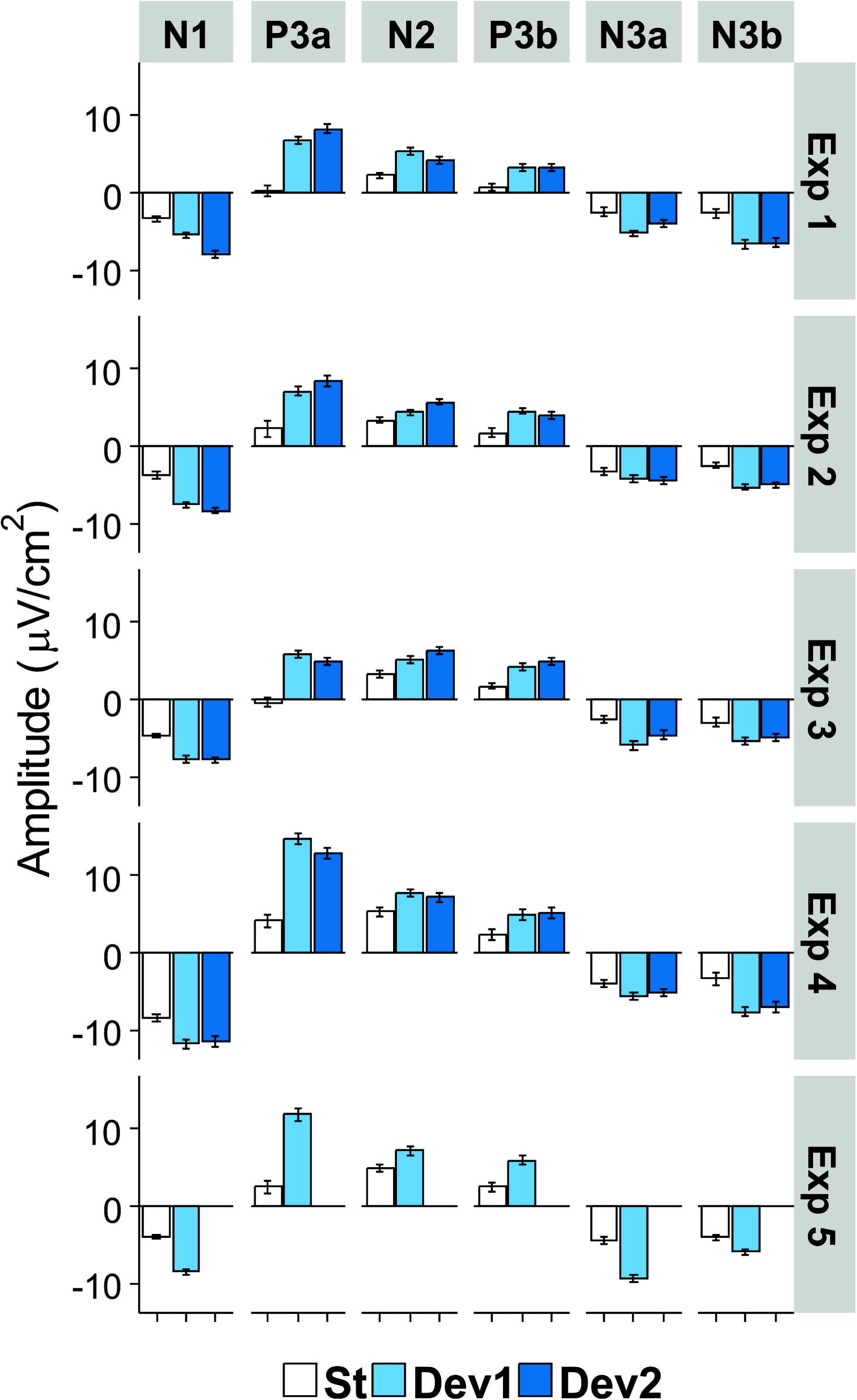
Mean amplitude of each ERP component in the region of its maximal expression (N1, frontocentral right; P3a, Cz; N2b, temporoparietal left; P3b, temporoparietal left; N3a (early) and N3b (late), frontocentral right). White bars, standard; light blue bars, Deviant 1 (or the only deviant in Experiment V); dark blue, Deviant 2. Bars show standard errors of means.

As regards **P3a**, both the mean component amplitude and its topographical differences were larger in Experiment IV than in the two other experiments (main effect of Experiment: F(2,36) = 21.24, p < .001, η^2^ = .54; Experiment x Site interaction: F(6,108) = 3.55, p = .022, η^2^ = .17). As regards **P3b**, the topographical differences between the lateral areas (i.e., Fro and FC negativities, Pos and TP positivities) were significantly smaller in Experiment II than in Experiments III and IV (Experiment x Area: F(8,144) = 4.33, p = .003, η^2^ = .19).

## Comparison between N1 and N3

### Methods

Given the apparent similarities in the distribution of N1 and N3, we conducted additional ANOVAs to find differences between these negative components. These ANOVAs were similar to those reported above but included an additional factor Principal Component (PC) having three levels (PC1, PC2, and PC4). Since absolute differences in magnitude between the components are not very informative, only interactions of PC with other factors will be reported.

### Results

The results were almost identical in all experiments. Highly significant interactions in lateral areas were obtained for the factors PC and Stimulus (Experiment I: F(4,88) = 13.57, η^2^ = .38; Experiment II: F(4,76) = 8.09, η^2^ = .30; Experiment III: F(4,80) = 12.81, η^2^ = .39; Experiment IV: F(4,84) = 8.69, η^2^ = .29; all p < .001), PC and Area (Experiment I: F(8,176) = 4.12, p = .022, η^2^ = .16; Experiment II: F(8,152) = 5.39, p = .002, η^2^ = .22; Experiment III: F(8,160) = 3.71, p = .035, η^2^ = .16; Experiment IV: F(8,168) = 22.64, p < .001, η^2^ = .52), as well as between PC, Stimulus, and Area (Experiment I: F(16,352) = 2.45, p = .039, η^2^ = .10; Experiment II: F(16,304) = 2.85, p = .002, η^2^ = .13; Experiment III: F(16,320) = 3.13, p = .007, η^2^ = .14; Experiment IV: F(16,336) = 2.36, p = .010, η^2^ = .10). An inspection of these interactions showed that the difference between standards and deviants was significantly larger for N1 than for both N3 subcomponents. Particularly, significant PC x Stimulus interactions (all p ≤ .002) were obtained in FC and Ce areas, where both the absolute negative values and the differences between standards and deviants were substantially larger for N1 than for N3. When PC1/N1 was excluded from the analysis (that is, the factor PC was taken with two levels), no effect of PC whatsoever approached significance.

Significant PC x Time interactions were obtained in Experiments II (F(4,76) = 5.39, p = .002, η^2^ = .22) and IV (F(4,84) = 4.31, p = .005, η^2^ = .17). The N1 amplitude significantly decreased with time (linear trend: F(1,21) = 8.25, p = .009, η^2^ = .28; and F(1,21) = 15.87, p = .001, for Experiments II and IV, respectively), but the amplitudes of N3 did not. Finally, Experiment IV showed a significant interaction between PC and Site *at midline* (F(6,126) = 5.40, p = .001, η^2^ = .20). While all negativities were characterized by the common trend to smaller negative (or larger positive) values in the posterior direction, the exact distribution was different: the negative-to-positive inversion took place already at Cz for N3, but only at Pz for N1.

## Experiment V

### Methods

Participants were 13 females and nine males, aged 19-33 (mean 25.0). Experiment V was, in fact, designed for a different purpose (investigation of affective conditioning), but included into the present report for a simple reason. Experiments I to IV all employed three stimuli and the similarity of their results might be attributed to this fact. In Experiment V, however, ERPs were recorded to two stimuli, and we wondered whether the same N3 effects would be obtained in this condition too.

This experiment also entailed an Acquisition phase and a Test phase. Five negative and 5 positive emotional exclamations were used. The stimuli and their effect on EEG and magnetoencephalographic (MEG) responses were reported in Kotchoubey et al. (2003), Bostanov and Kotchoubey (2004), and Kotchoubey et al. (2009). Emotionally positive and negative exclamations did not differ from each other in terms of the average duration and fundamental frequency (F0), and their average loudness was equalized as well. The Acquisition phase consisted of 50 presentations of Chord 1 paired with five positive exclamations (10 times each exclamation) and 50 presentations of Chord 2 paired with five negative exclamations (10 times each). The two chords were 440 + 880 + 1760 + 3520 + 7040 Hz and 150 + 300 + 600 + 1200 + 2400 Hz, and their linkage to positive and negative emotional exclamations was counterbalanced among participants. Tone duration was 200 ms, SOA between the tone and the exclamation 300 ms, and SOA between pairs randomly varied from 2000 to 2280 ms. The following Test phase was an oddball paradigm: one of the chords presented during Acquisition was randomly selected as the standard (presented 300 times), and the other one, as the deviant (presented 60 times). Emotional exclamations were not presented in the Test phase anymore. SOA in the Test phase randomly varied between 950 and 1050 ms.

The same PCA procedure was applied to Experiment V data separately from the other four experiments. 44 temporal factors were extracted for rotation. Despite numerous methodological differences between the experiments, the PCA resulted in surprisingly similar distribution of the main components (see Figure 7). The only major difference was the presence of PC7 (peak latency 376 ms) that could not be identified as a peak in the ERP waveform. As shown in Table 4, other PCs were almost identical to those in Experiments I – IV, only their order was changed.

**Figure 7.**
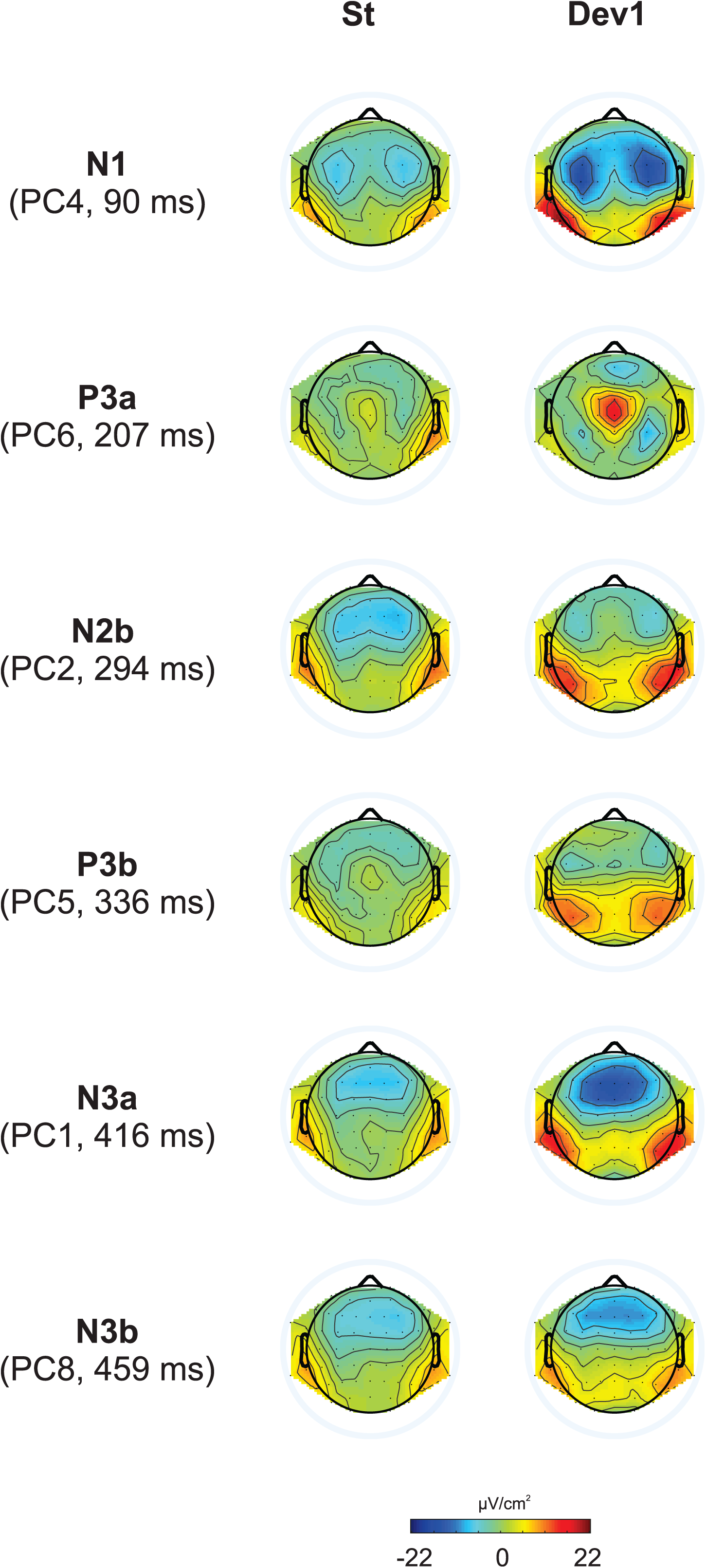
Topographic representation of each principal component at their peak latencies for Experiment V.

**Table 4.** Principal components selected in the experiments with three stimuli (Experiment I to IV) and in Experiment V with two stimuli. Note very close peak latencies of the PCs obtained with different stimulation

| Peak name |  | N1 | P2 | P3a | N2b | P3b | N3early | N3late |
| --- | --- | --- | --- | --- | --- | --- | --- | --- |
| Exp I-IV | Principal component | PC1 | PC6 | PC5 | PC3 | PC7 | PC2 | PC4 |
|  | Latency, ms | 96 | 136 | 215 | 294 | 334 | 412 | 456 |
| Exp V | Principal component | PC4 | PC3 | PC6 | PC2 | PC5 | PC1 | PC8 |
|  | Latency, ms | 90 | 129 | 207 | 294 | 336 | 416 | 459 |

### Results

The main object of the present study, **N3**, was again represented as two subcomponents, PC1 and PC8. At midline, both were negative at frontal and central sites but positive at Pz (main Site effect). Both fronto-central negativities and parietal positivities were about twice larger to deviants than to standards (Stimulus x Site interactions). The analysis over lateral sites revealed negative amplitudes in Fro and FC areas, and positive amplitudes in Pos and TP areas (main effect of Area). The topographical differences were larger for deviants than standards (Table 5), resulting in highly significant Stimulus x Area interactions. The amplitude of PC1 (but not PC8) appeared to decrease with time, particularly at frontal sites on midline and lateral Fro and FC areas, but neither the main effect of Time, nor the Time x Site and Time x Area interaction attained the corrected significance level.

**Table 5.** Anterior-posterior differences in the amplitudes (in µV/cm^2^) of two Principal Components (PCs) reflecting N3. Fro – frontal, TP – temporoparietal. “Lateral” means averaged values for the left and right hemispheres, because the factor Hemisphere was not significant.

|  |  | Fro midline | Pz | Fro lateral | TP lateral |
| --- | --- | --- | --- | --- | --- |
| PC1 | standard | -2.9 | -9.8 | -4.2 | -11.1 |
|  | deviant | 1 | 4.8 | 4.7 | 4.4 |
| PC8 | standard | -2.7 | -5.3 | -4.8 | -6 |
|  | deviant | 1.9 | 3.7 | 2.9 | 5.1 |

**N1** (**PC4)** had its midline maximum at frontal and fronto-central sites with a decreasing amplitude toward Pz. The amplitude to deviants was larger than to standards at all sites except Pz, resulting in a significant interaction between Stimulus and Site. At lateral sites, the amplitude was negative (and more negative to deviants than standards) in the Fro and FC areas, and positive (and more positive to deviants than standards) in the TP areas, leading to strong main effects of Area, Stimulus and their interaction.

**N2b (PC2)** was positive related to the baseline at Cz, CPz, and Pz and close to zero frontally. At lateral sites its amplitude became more positive in the posterior direction. Also, the amplitude was generally positive to deviants and negative to standards.

Both positive components, **P3a (PC6)** and **P3b (PC5)**, decreased at midline with Time and were more positive to deviants than standards. The stimulus effect on P3a was best expressed at Cz, and the stimulus effect on P3b, at Pz (but Stimulus x Site interaction reached significance only for P3a).

The lateral distribution of the positive components strongly differed between standards and deviants. Although both stimuli elicited negative **P3a** amplitudes in Fro areas, the maximum positivity to standard was observed in TP areas, but the maximum positivity to deviant was much more anterior, in FC areas. **P3b** amplitudes to standards were slightly negative in almost all areas (Fro, FC, Ce, Pos), but positive in TP areas, whereas P3b amplitudes to deviants were negative in FC areas, zero in Fro areas, and positive in Ce, Pos and TP areas. Thus both components displayed the main Area effect and the Stimulus x Area interaction. Averaged across all areas, mean amplitudes were positive to deviants and slightly negative to standards.

Like in the previous experiments, we directly compared N1 with N3 in Experiment V because superficially they appear to display similar dynamics. PC1 was taken as the best representative of N3, and PC4, as the representative of N1. Thus the factor PC had 2 levels.

In the lateral ANOVA the factor PC strongly interacted with Stimulus (F(1,21) = 30.13, p <. 001, η^2^ = .59) and Area (F(4,84) = 9.09, p <. 001, η^2^ = .30), also yielding a three way PC x Stimulus x Area interaction (F(4,84) = 4.71, p =. 006, η^2^ = .18). Generally, the difference between standards and deviants was significantly larger for PC4/N1 than for PC1/N3. The topographic distributions were different: N1 was negative in the Fro, FC, and Ce areas, zero in the Pos areas, and positive only in the TP areas, and the mean difference between the most negative Fro areas and the most positive TP areas was 8.8 µV/cm^2^. N3 was negative in the Fro and FC areas, zero in the Ce areas, and positive in the Pos and TP areas, and the difference between the most negative and most positive areas was 12.3 µV/cm^2^. The maximal effect of stimulus (> 4 µV/cm^2^ difference between deviant and standard) was obtained in the Ce areas for N1 but in the FC and TP areas for N3.

The analysis at midline replicated the interactions with topography (PC x Site: F(3,63) = 8.19, p =. 001, η^2^ = .28; PC x Stimulus x Site: F(3,63) = 4.84, p =. 016, η^2^ = .19). Like at the lateral sites described above, the transition from negative to positive values happened more posteriorly for PC1/N3 than for PC4/N1, and the difference between maximal negative and maximal positive values was larger for the former than the latter (5.1 and 8.1 µV/cm^2^ for PC4/N1 and PC1/N3, respectively). The largest effect of stimulus was at Cz for PC4/N1 but at frontal leads for PC1/N3. Figure 8 presents the comparison between N3 and N1 in three experiments.

**Figure 8.**
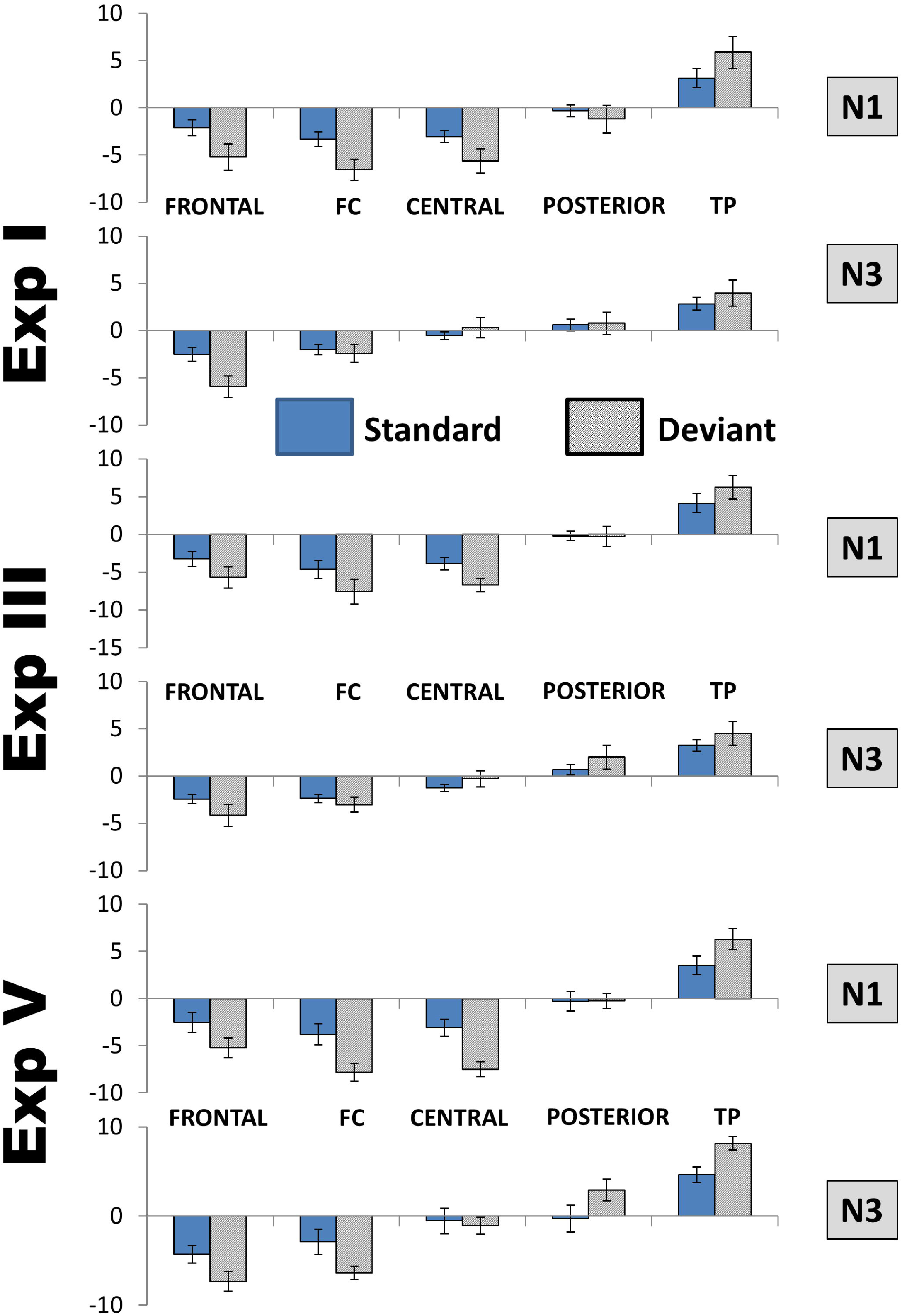
Comparison between the distributions of N1 and N3 over lateral areas. FC, frontocentral areas; TP, temporoparietal areas. To save space only the data of three experiments are presented, but the distributions were not qualitatively different also in the other two. Because of only minimal hemispheric differences, left and right areas are averaged, as well as Deviant1 and Deviant 2 in Experiments with two deviants. Amplitudes are in µV/cm^2^. Bars show standard errors of means.

For control, this analysis was repeated using PC8 instead of PC2. According to our inspection, PC8 reflected a later subcomponent of the same component N3. In fact the analysis of PC1 versus PC8 did not reveal any significant interactions of the factor PC.

### Discussion

The analyses of Experiments II, III, and IV together, as well as the results of Experiment V, confirm the stability of the N3 component and its relative independence of the design of oddball experiments. The effects of the methodical differences between the experiments were very weak if any. The data further dissolve the view of the apparent similarity between N3 and N1: when the two components are included in an ANOVA as a factor, substantial differences between them are revealed. In contrast, when two N3 subcomponents are included in an ANOVA as a factor, this factor does not interact with any other indicating that the two subcomponents reflect basically the same thing.

## Mastoid references

Because many studies standardly report ERP data using average mastoid reference, we additionally carried out the same PCA for mastoid-referenced data to enhance comparability with the literature. The results were very similar for all experiments. Therefore, here we report the results of an analysis including the data of all five experiments together. In the experiments with two deviant stimuli, responses to all deviants were averaged together to obtain one standard and one deviant waveform. The PCA yielded a total of 41 Factors. The following analysis was performed like for CSD data.

As can be seen in Figure 9, principal component peaks had latencies very similar to CSD-extracted components: 96 ms for N1 (PC1), 215 ms for P3a (PC5), 334 ms for P3b (PC7), 412 and 456 ms for two subcomponents of N3 (PC6 and PC9, respectively). The main difference between CSD and mastoid referenced N3 is that in the latter case, N3 decreased in the posterior direction but did not reverse its polarity; that is, it remains negative at all sites.

**Figure 9.**
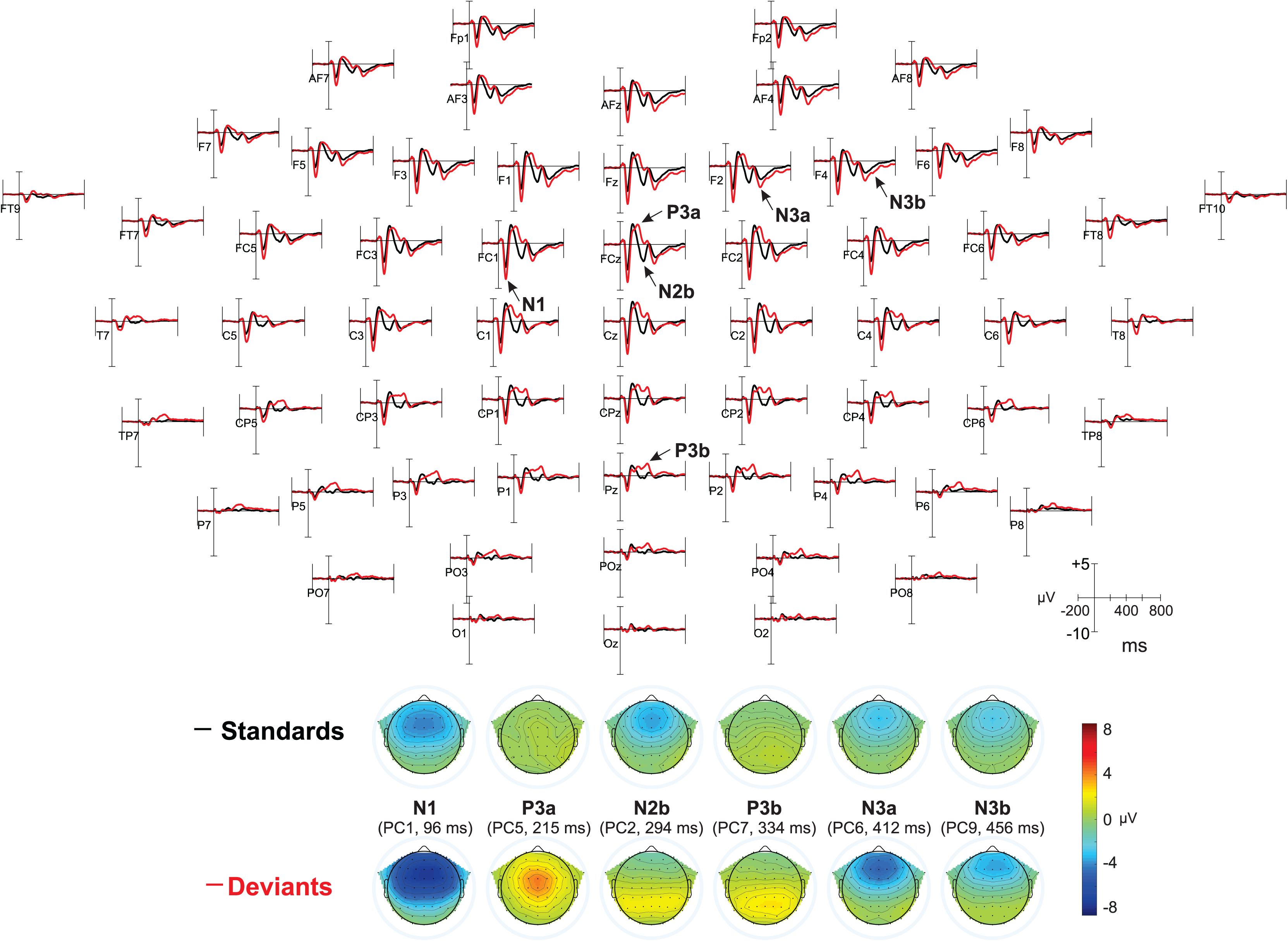
Results of mastoid-referenced data. Top: ERP waveforms averaged for all five experiments, referred to average mastoids, for standards (black) and deviants (red). In Experiments I – IV two deviants were taken together. Most prominent peaks are indicated by arrows (P2, which did not differ between standards and deviants, is not shown). Bottom: topographical distributions for PCA data.

We omit the results of the statistical analysis of mastoid-referenced data (which was performed in exactly the same manner as for CSD data) because this would considerably increase the volume of the article without any added information value.

## General Discussion

### Reliability

To our best knowledge, this is the first systematic investigation of the N3 wave in the oddball paradigm. Being the first, it is largely exploratory in character. We conducted a large number of analyses, some of which with up to five factors. Such analyses can yield false positive results. We are aware both of this problem and of the fact that it still does not have a satisfactory solution. Thus we admit that some effects described in the Results section may later turn out to be spurious. Nevertheless, we have several reasons to be convinced that at least the most important findings are reliable. First, many of them attained very high significance level with p ≤ .001, which would survive any reasonable correction. But the second reason for our optimism is much more important than the formal level of significance: the major effects are consistently *replicated* from experiment to experiment notwithstanding considerable differences between experimental conditions. This pleasantly surprising replicability already starts with the results of PCA. The latencies of principal components found in different experiments differed, on average, in 4.8 ms; the latencies of the components reflecting N3 (the target component of the present study) differed in 2.7 ms only. In other words, the structure of ERP was virtually identical in all experiments.

N3 is obviously not an artifact. In our experiments all kinds of artifacts were carefully controlled. Participants listened to stimuli with eyes closed, which strongly diminished ocular effects. Bad channels were replaced by interpolation, non-EEG components were removed by means of the ICA, and finally, a few remaining artifact trials were excluded from averaging. Also in the literature we could find no indication of the artificial nature of N3.

The PCA showed that N3 consistently entailed two subcomponents only weakly correlating with each other, with peak latencies of 412-416 and 456-459 ms, respectively. All experimental effects, however, were virtually the same on the two subcomponents and revealed only quantitative differences (mostly, the earlier subcomponent exhibited stronger effects than the later one). Both subcomponents also had the same topography.

### Differentiation from other phenomena

The strong similarity between the earlier and the later N3 subcomponents can be compared with the significant differences between them and the other ERP negativities. Although a superficial glance might see a similar distribution of N3 and N1 (negativities in frontal areas, positivities in temporo-parietal areas), their direct comparative analysis demonstrated substantial differences. The point of the negative-to-positive inversion was much more anterior for N3 than for N1. The maximum of frontal negativity was more lateralized for N1 than N3. The distribution of N1 CSD is compatible with a fronto-temporal sink, whereas the distribution of N3 CSD is rather in line with a midfrontal sink. Further, N1 demonstrated a tendency to habituation, but N3 did not.

As all stimuli in our experiments had a duration of 200 ms, one might even ask whether N3 was an N1 of the *offset* response. But this hypothesis is, of course, wrong just because other studies (including our own: Kotchoubey and Pavlov, 2017) used stimuli of broadly varying duration, yet the peak latency of N3, according to the corresponding figures, was always between 400 and 450 ms from stimulus *onset*. Likewise, N3 is obviously not a mismatch negativity. CSD data of the MMN demonstrate a completely different topographical distribution (see, for example, Deouell et al., 1998, Fig. 3b; Shalgi and Deouell, 2007, Fig. 5; Kayser and Tenke, 2015, Fig. 12). Furthermore, the number of stimuli in the present experiments was rather small. N3 could be seen even after as few deviants as 20-50. Although a few studies showed the MMN after 50-70 deviants, most typical MMN experiments use hundreds of deviants to demonstrate consistent effects.

The decrease of the N3 amplitude in the posterior direction might remind on the Positive Slow Wave (PSW, e.g., Squires et al., 1975; Sutton and Ruchkin, 1984; Dien, 1998; Spencer et al., 1999; Dien et al., 2004) that can also have negative amplitudes in frontal leads. However, the analysis of the data with average mastoid reference (Figure 8) shows that, contrary to the PSW, N3 does not change its polarity in the sagittal dimension, i.e., it remains negative also in the parietal areas. This is in line with CSD data indicating that the electrical source of the N3 dipole is not parietal but posterior temporal, in the vicinity of the mastoid electrodes. Furthermore, Kayser et al. (1998) identified N3 and the PSW as two different principal components.

Other two late components that can be mentioned *post hoc* are the O-wave (Näätänen et al., 1982) and the reorienting negativity (RON: Schröger & Wolff, 1999; Berti, 2008). We did not think on these components in advance because the conditions of our experiments have nothing in common with those in which O-wave and the RON are ordinarily obtained. The former occurs in attended conditions, usually when a motor response is required to the next expected stimulus. It is long lasting and can (like the PSW above) be inverted at parietal sites, while N3 is recorded in the passive condition, relatively short-lasting and always negative except over posterior temporal regions.

The RON, as one of its discoverers emphasized, “was observed after distraction *only in situations when a reorientation on the task at hand was required*”; therefore, “the RON was interpreted as the correlate of a switch of attention onto the task-relevant information” (Berti, 2008; p.609; italics added). Most usually, a RON follows P3a without a P3b, contrary to N3 that follows P3b. Sometimes, however, the positivity prior to a RON contains two components called “early P3a” and “late P3a” (e.g., Correa-Jaraba et al., 2016, 2018), and there is no strong argument as of why the latter cannot, in fact, be a P3b. In some exceptional cases one can see a clear P3b before the RON (e.g., Scheer et al., 2016).

In respect of time course and morphology, the RON is much more similar to the passive N3 than any other late components such as O-wave or PSW. Moreover, a few studies using CSD analysis also indicate a surprising similarity in the topography (e.g., Rämä et al., 2018). To sum up, there is an astonishing contradiction between the obvious similarities of N3 and the RON, on the one hand, and the drastic differences in the experimental conditions in which these two waves are elicited. The contradiction can, from our point of view, be solved in one of two ways. Either a more comprehensive analysis will detect substantial differences between the RON and N3 as two independent components, or, if they are identical, the present-day interpretation of RON (including its processing specificity) will be seriously challenged, because the typical conditions of N3 do not involve any reorientation process.

Furthermore, N3 does not appear to depend on the alleged significance of stimuli, at least within the range tested in this study. Tones, which elicited N3, had been previously related to participants’ own names in Experiments I and IV, to emotional stimuli in Experiment V, to unconceivable sounds in Experiment III, and were completely neutral in Experiment II. No significant differences between the results of these experiments were observed. Likewise, N3 did not substantially depend on the number of stimuli in the oddball experiment, as it was equally recorded in two- and three-stimuli paradigms.

Finally, N3 demonstrated an oddball effect, being larger to deviants than standards. In contrast to N1 and P3, this effect was rather manifested as a Stimulus x Topography interaction than as a main effect of Stimulus. This means that N1 was consistently more negative, and P3a and P3b were more positive to deviants than standards, although these amplitude increases also had their local maxima for each component. N3, in contrast, was typically more negative to deviants than standards over anterior regions but less negative to deviants than standards over posterior regions.

### Passive N3 and its hypothetical functional meaning

Table 5 shows an overview of the presence of N3 in the previously published passive oddball reports. The search was performed on PUBMED using the terms “passive oddball” or “passive P3”. Because we assume that N3 is a sharp negativity following P3b, we included only studies whose results indicate a P3b (and not only the complex MMN-P3a). Another inclusion criterion was that ERP waveforms at least at the three midline sites (Fz, Cz, Pz) were clearly presented.

**Table 6.**
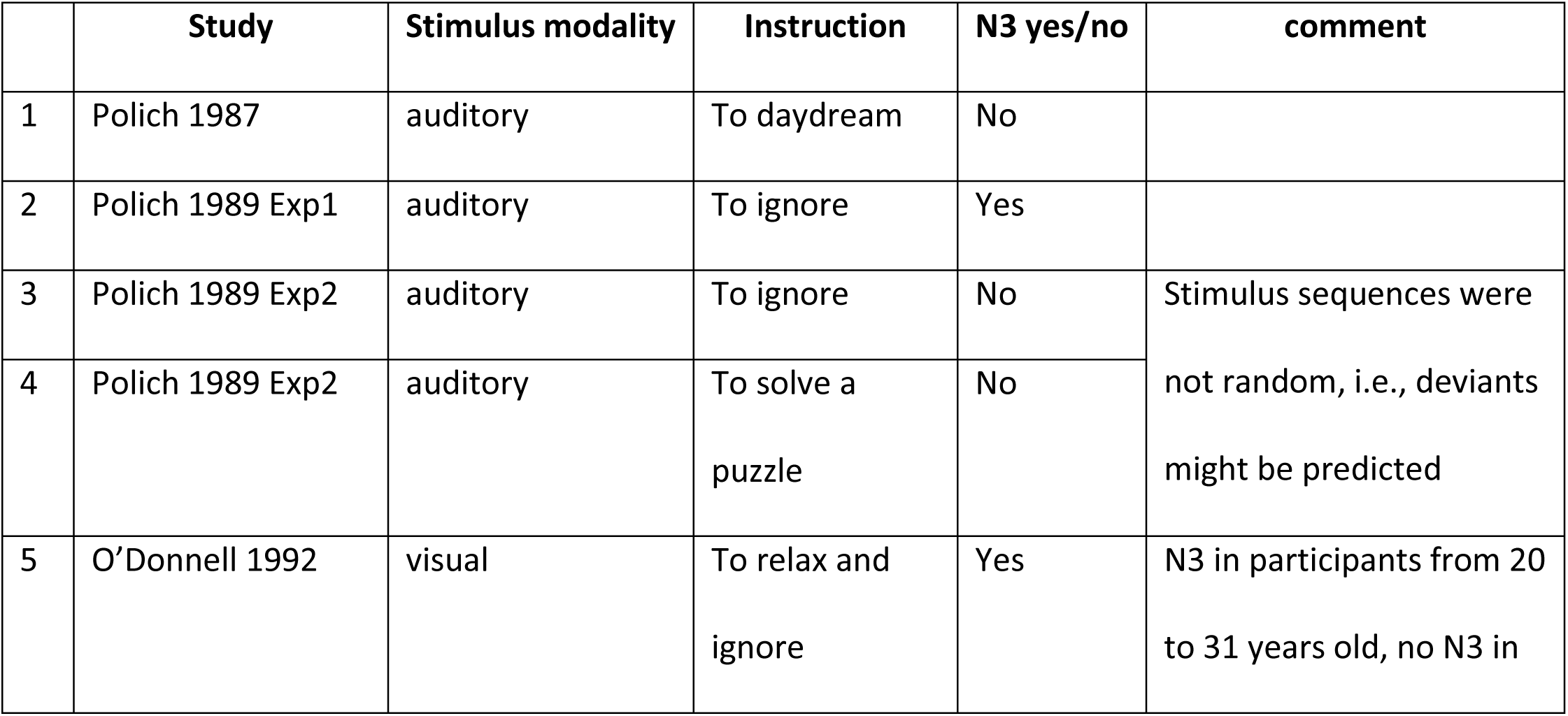
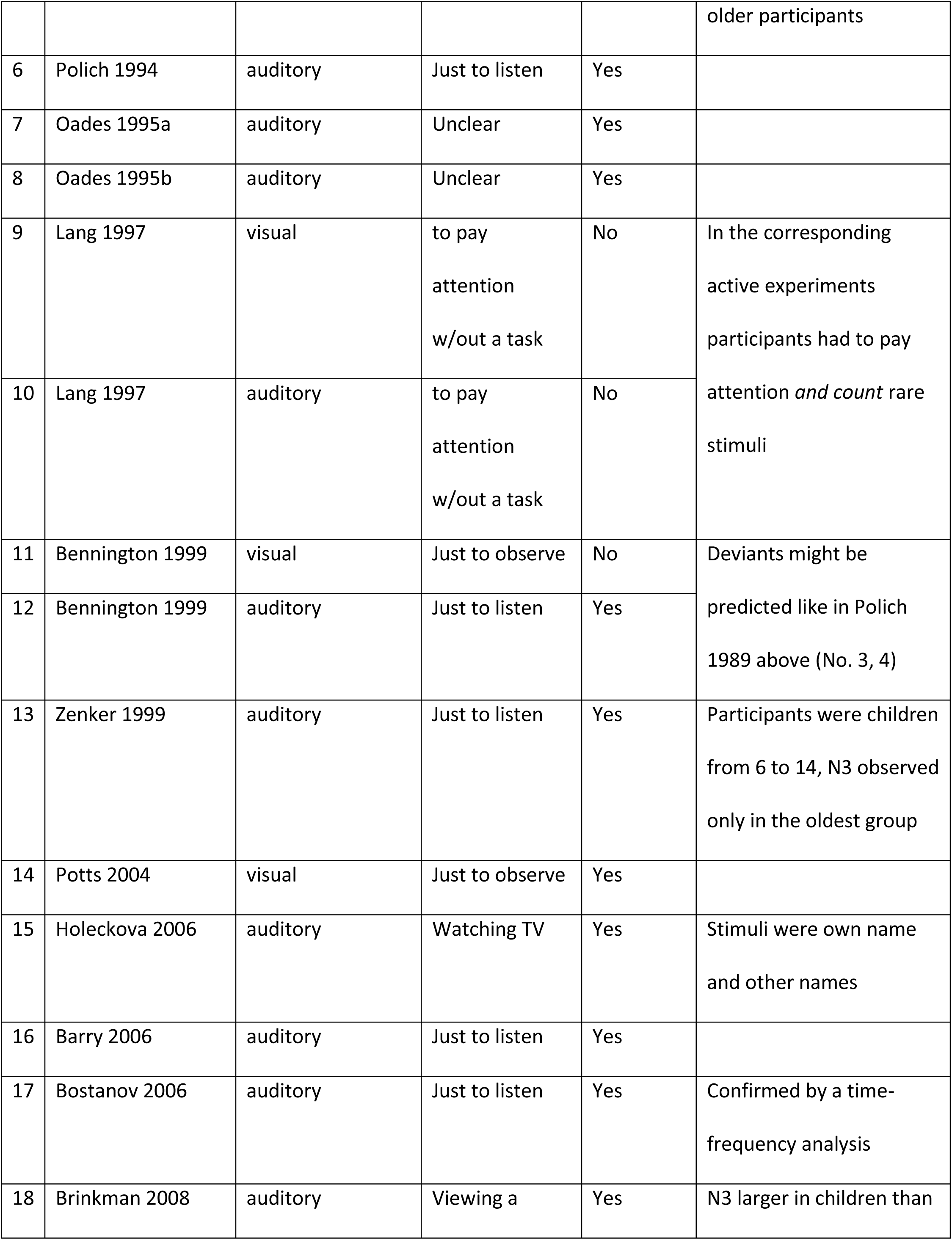
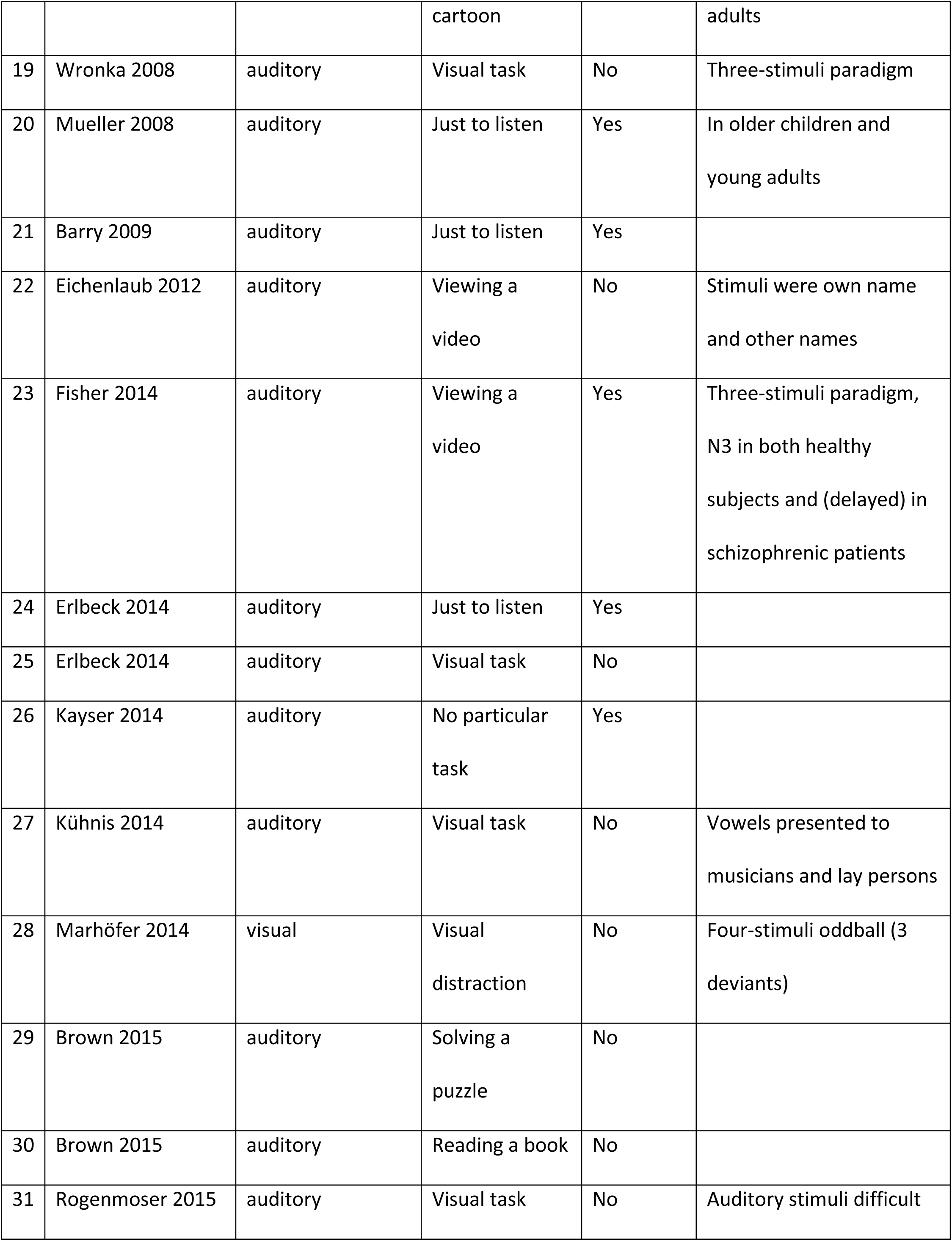
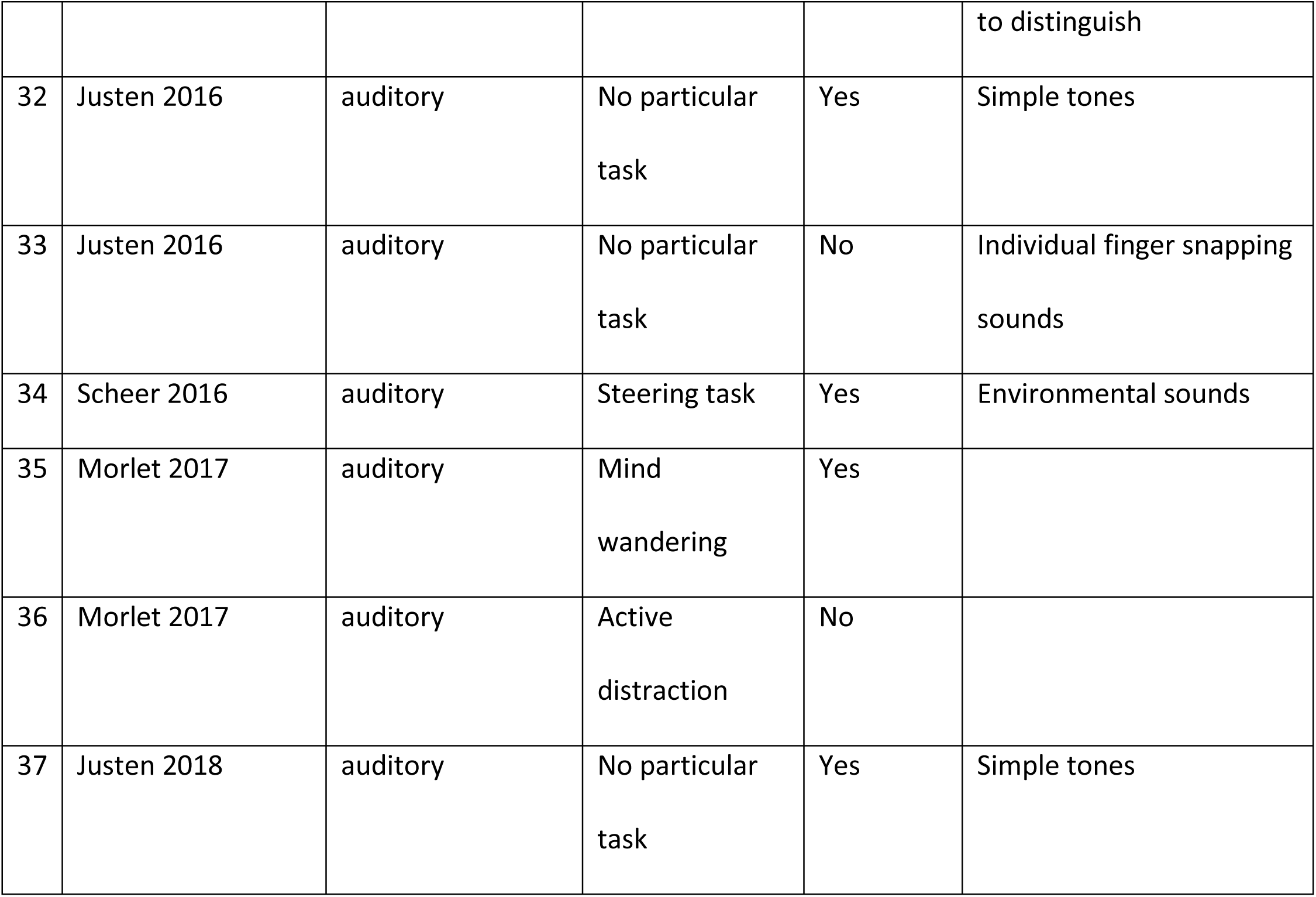
Experiments designated as “passive oddballs”, in which figures do or do not show an N3 following P3b.

References to the table: (Barry et al., 2006; Barry et al., 2009; Bennington and Pollich, 1999; Bostanov and Kotchoubey, 2006; Brinkman and Stauder, 2008; Brown et al., 2015; Eichenlaub et al., 2012; Erlbeck et al., 2014; Fisher et al., 2014; Holeckova et al., 2006; Justen and Herbert, 2016; Justen and Herbert, 2018; Kayser et al., 1997; Kühnis et al., 2014; Lang et al., 1997; Marhöfer et al., 2014; Morlet et al., 2017; O’Donnell et al., 1997; Oades et al., 1995a; Oades et al., 1995b; Polich, 1987; Polich, 1989; Polich and McIsaak, 1994; Potts, 2004; Rogenmoser et al., 2015; Scheer et al., 2016; Wronka et al., 2008; Zenker and Barajas, 1999).

The table lists a total of thirty-seven experimental conditions, N3 being observed in twenty-one of them. Of these 21, the condition described as “passive” included an additional distraction task in four experiments: viewing a cartoon in Brinkman and Stauder (2008), viewing a video in Holeckova et al. (2006) and Fisher et al. (2014), and a demanding steering task in Scheer et al. (2016). Inspecting these four experiments, one can see that Holeckova et al. (2006) presented participants’ own names and similar names as the to-be-ignored auditory stimuli, and one may suppose that the distraction task did not completely distract participants from the processing of the highly significant words. In Scheer et al. (2016) a strong N3 after P3b was elicited by highly attractive environmental stimuli such as laughing, but a rather small N3 (without a preceding P3b) was also observed in response to tones. Participants of Brinkman & Stauder (2008) were adolescents, and those of Fisher et al. (2014), schizophrenic patients. However, in the latter study N3 was also observed in the control group. In the other 17 experiments showing N3, the authors implicitly (Oades et al., 1995a, b) or explicitly (the rest of studies) indicated the lack of any task: participants should simply view visual stimuli or listen to auditory ones.

Of the 16 data sets that do *not* show an N3, active distraction was used in nine experiments. Moreover, the participants in both visual and auditory experiments of Lang et al. (1997) were instructed to actively follow stimuli, albeit without a motor task. Polich (1987) instructed his subjects to daydream, which may or may not be regarded as distraction. In his later works (Polich, 1989) a passive condition without any distraction was presented twice, and N3 was recorded in Experiment 1 but not in Experiment 2. The sequence of auditory stimuli in this study and in Bennington and Polich (1999) was not random or pseudorandom, but consisted of ten-stimuli trains, and deviants could only appear in the second halves of the trains. This means that subjects could guess deviants more or less successfully. In an extreme case, after a series of 9 standards, the deviant could even be predicted with certainty. Finally, Justen and Herbert (2016) showed an extremely large N3 in a completely passive condition with tonal stimuli, but very small (if any) in the same oddball with sounds of snapping fingers. Again, such sounds (also including snapping of a participant’s own fingers) are highly idiosyncratic.

Of course, passive conditions present a principal psychological problem, because the less exact is participants’ instruction, the less do we really know what they were doing during stimulation. This general difficulty is also valid for the experiments summarized in Table 5. However, the above analysis results in a rough differentiation between the two kinds of conditions that can be designated as “passive-attentive”, in which participants’ attention was directed either to a different class of stimuli or to the presented stimuli (but without any response requirement), and “passive-inattentive”, in which participants’ attention was allowed to freely move around. If this classification is correct, N3 was shown in 17 “passive-inattentive” conditions and 4 “passive-attentive” conditions. No N3 was shown in 5 “passive-inattentive” conditions and 11 “passive-attentive” conditions. The resulting difference is highly significant (Fisher Exact Test: p = .006).

This leads to a hypothesis that N3 may be a result of an intermediate level of stimulus processing that differs from both very superficial and very deep processing. Generally, slow cortical negativities reflect preparatory activation of apical dendrites, and thus anticipatory “warming up” large populations of cortical neurons (Rockstroh et al., 1989; Birbaumer et al., 1990; Mitzdorf, 1991). Because such preparation to a further activity is usually followed by the phase of the realization of the prepared potential, the vast majority of cortical ERPs can be presented as a sequence of negative-positive complexes (i.e., cycles of preparation and realization: Kotchoubey, 2006). In some cases, however, a cortical response ends with a negative rather than positive phase, such as the occipital P1-N1 complex of visual ERPs or the N400 to semantically unrelated word pairs (e.g., *table-mouse*) presented on the background of related word pairs (e.g., *cat-mouse*). Such an ending negativity is usually a sign of preparation that did not find its realization (Kotchoubey, 2006).

According to this hypothesis, N3 should be elicited neither during a goal-directed activity, when the eliciting stimulus is regarded as task relevant, and the corresponding task is performed, nor in a situation of attentional distraction when the stimulus is regarded as fully irrelevant and processing resources are directed to a completely different task. In the former case, there are two options. With simple task-relevant stimuli, such as in most oddball experiments, the processing is closed with a P3b whose meaning has been a subject of long discussions (e.g., Johnson, 1986; Donchin and Coles, 1988; Verleger, 1988; Verleger et al., 1994; Kotchoubey et al., 1997). Very complex or highly emotional stimuli elicit, beyond this, late slow wave activity that reflects sustaining processing. In the latter case again two options appear: if an irrelevant stimulus matches the current expectancies, its processing ends with the P2 component some 180-250 ms after stimulus onset; if it mismatches the expectancies, the complex of the MMN and P3a follows. None of these four cases requires an additional negative deflection N3 after P3b.

The situation is different when the stimulus is regarded by the brain as potentially relevant, yet its supposed relevance cannot be linked to any adaptive activity. It appears to mean something, but no behavioral consequence can be inferred from this meaning. The result is processing that commences but does not come to a closure.

If this hypothesis is true, further experiments can be designed to vary the precision of the meaning of presented stimuli. A starting paradigm may be the design of Erlbeck et al. (2014) or Morlet et al. (2017) who employed an active condition (attention to the stimuli), a distraction condition (attention away from the stimuli), and a passive condition (free attention). However, future experiments on this basis should employ more gradual conditions, such as attentional tasks with high and low motivation (assuming that motivation mobilizes attentional resources), instructions to pay attention to the stimuli without any overt or covert response, or just presentation of stimuli without any instruction. Also, though the results of the above studies *appear* to be in line with the hypothesis (i.e., a manifested N3 is obtained in the passive condition only), nothing can be said with confidence because the authors were primarily interested in earlier components such as the MMN and P3.

Moreover, if this hypothesis is true, N3 might also be found in some *active* oddball experiments, e.g., when the experimental instruction was (intentionally or not) formulated with less precision. For example, N3 in active oddball was recorded in Polich (1989) and Kayser et al. (1998), and in Müller et al. (2008) it was even larger in the active than passive paradigm, which is in strong contrast with the data of Erlbeck et al. (2014), Brown et al. (2015) and Morlet et al. (2017). However, the hypothesis about a link between the active N3 and the precision of the experimental instruction can only be tested in a systematic review of the voluminous literature about ERPs in active oddball, which should be a subject of a separate study.

## Acknowledgement

The authors thank Janna Holst and Ines Armbruster for their help in carrying out the experiments.

